# A permeable protein nanocage enables facile cargo loading and cytosolic protein delivery

**DOI:** 10.64898/2026.04.06.716810

**Authors:** Seokmu Kwon, Michael P. Andreas, Jesse A. Jones, Tobias W. Giessen

## Abstract

The cytosolic delivery of therapeutic proteins remains one of the most persistent challenges in modern drug delivery. Here, we report the discovery and characterization of an encapsulin-based protein nanocage, QtEnc, with unexpected permeability properties and the ability to internalize cargo proteins *in vitro*, fundamentally departing from existing protein nanocage cargo loading paradigms. This permeability enables simple, rapid, and single-step post-assembly cargo loading, accommodating cargos as large as 482 kDa, and allowing multiplexed cargo co-encapsulation with tunable ratios. Leveraging this property, we develop a modular QtEnc-based NanoCarrier (QtEncNC) with a pH-responsive cargo detachment module and an endosomal escape module, enabling low pH-triggered cargo release from assembled shells and subsequent endosomal escape for cytosolic delivery. Using a cytotoxic protein, BLF1, as a proof-of-concept QtEncNC payload, we demonstrate efficient cytosolic protein delivery in HeLa cells. These findings establish QtEncNC as a versatile and modular platform for cytosolic protein delivery with broad biomedical potential.

---

Nanocarrier-based drug delivery systems represent an innovative approach in modern medicine for the targeted delivery and controlled release of therapeutics.^1,2^ Biologics in particular can benefit from the protection,^3^ targeting ability,^4^ increase in bioavailability,^5^ and enhanced circulation times^6^ provided by different nanocarrier platforms. Nanocarriers have the potential to enhance the efficacy and safety of drugs while decreasing side effects.^1,7^ Whereas some nanocarrier-based drugs can be successfully aimed at extracellular targets,^8^ accomplishing cytosolic delivery of especially therapeutic proteins has remained challenging.^9^ As the cellular uptake of many nanocarriers is mediated by endocytic pathways,^10^ achieving triggered and appropriately timed drug release and endosomal escape represent formidable challenges. Considering that the majority of therapeutic targets are found within cells,^11^ designing nanocarriers capable of efficient cytosolic delivery has the potential to provide innovative solutions to many challenges in cancer therapy,^12^ enzyme replacement therapy,^13^ immunotherapy,^14^ gene editing,^15^ and vaccine development.^16^

The inherent difficulty of cytosolic drug delivery makes the continued discovery, design and engineering of novel potential nanocarrier systems an important strategy to advance the field. One such system are encapsulins, a fairly recently discovered class of natively protein-loaded prokaryotic nanocompartments.^17^ Encapsulin shell proteins self-assemble into icosahedral nanocages ranging in size from ca. 20 to 45 nm in diameter with triangulation numbers of T = 1 (60 subunits), T = 3 (180 subunits), or T = 4 (240 subunits).^18,19^ Their eponymous feature is the ability to specifically encapsulate dedicated cargo proteins *in vivo*.^20^ All native cargos contain cargo loading peptides (CLPs) that mediate encapsulation during shell self-assembly inside prokaryotic cells.^21^ As CLPs are highly modular, they have been utilized to package non-native cargo proteins into encapsulin shells by genetically fusing CLPs to proteins of interest.^22–26^ Importantly for engineering applications, encapsulin shells are often highly robust and have been modified through genetic insertions^27,28^ or fusions^29,30^ and chemical conjugation^31^ without disrupting cargo loading or shell assembly. While various ways of conferring targeting capabilities to encapsulins have been explored and applied in cell culture settings,^28,32,33^ cytosolic cargo delivery—especially of proteins—has remained challenging. This is likely due to the difficulty of engineering shells to undergo stimulus-triggered disassembly, achieving efficient cargo−shell detachment, and enabling endosomal escape of the released cargo within a single nanocage platform.

Here, we report the discovery and characterization of a permeable encapsulin-based protein nanocage, QtEnc,^34^ and its application as a nanocarrier for cytosolic protein delivery. We highlight QtEnc’s ability to efficiently internalize diverse cargo proteins *in vitro*, an unprecedented encapsulin property that enables simple post-shell assembly cargo loading that fundamentally differs from existing protein nanocage cargo loading paradigms. We further demonstrate that this *in vitro* loading mode can accommodate a wide range of cargo sizes and allows multiplexed cargo loading. Building on these properties, we rationally designed a modular QtEnc-based NanoCarrier (QtEncNC) for cytosolic protein delivery, which exhibits efficient cellular uptake, promotes endosome-specific cargo release without requiring shell disassembly, and ultimately achieves cytosolic protein delivery. Together, these findings establish QtEncNC as a versatile and modular platform with broad potential across diverse biomedical application areas, including cancer therapy, enzyme replacement therapy, and beyond.

## QtEnc shells can internalize proteins *in vitro*

The QtEnc iron storage encapsulin system is encoded by a three-gene operon found in *Quasibacillus thermotolerans* and consists of the encapsulin shell protein QtEnc, the primary ferroxidase cargo protein IMEF, and a minor 2Fe-2S ferredoxin cargo (Fdx) (Fig. 1a). The QtEnc shell self-assembles into a 240 subunit 42 nm icosahedral shell (T = 4) with one internal CLP binding site per subunit. It has previously been shown that IMEF-loaded QtEnc shells can sequester large amounts of internalized ferric iron in the form of mineralized ferrihydrite, playing an important role in cellular iron storage and homeostasis.^34^ The function of Fdx is still unknown but in analogy to the ferredoxin-mediated iron release from bacterioferritin cages,^35^ a role of Fdx in transferring electrons into the QtEnc shell has been proposed. This would allow the reduction, solubilization, and re-mobilization of stored iron. While IMEF possesses a C-terminal CLP, Fdx is the only known encapsulin cargo carrying a CLP at its N-terminus (Fig. 1a). Both CLPs possess identical core CLP motifs (TVGSL) crucial for cargo loading. Fdx is composed of a 39-residues long disordered N-terminal extension—within which the CLP core motif resides—and a C-terminal globular 2Fe-2S domain (Fig. 1b).

**Fig. 1.**
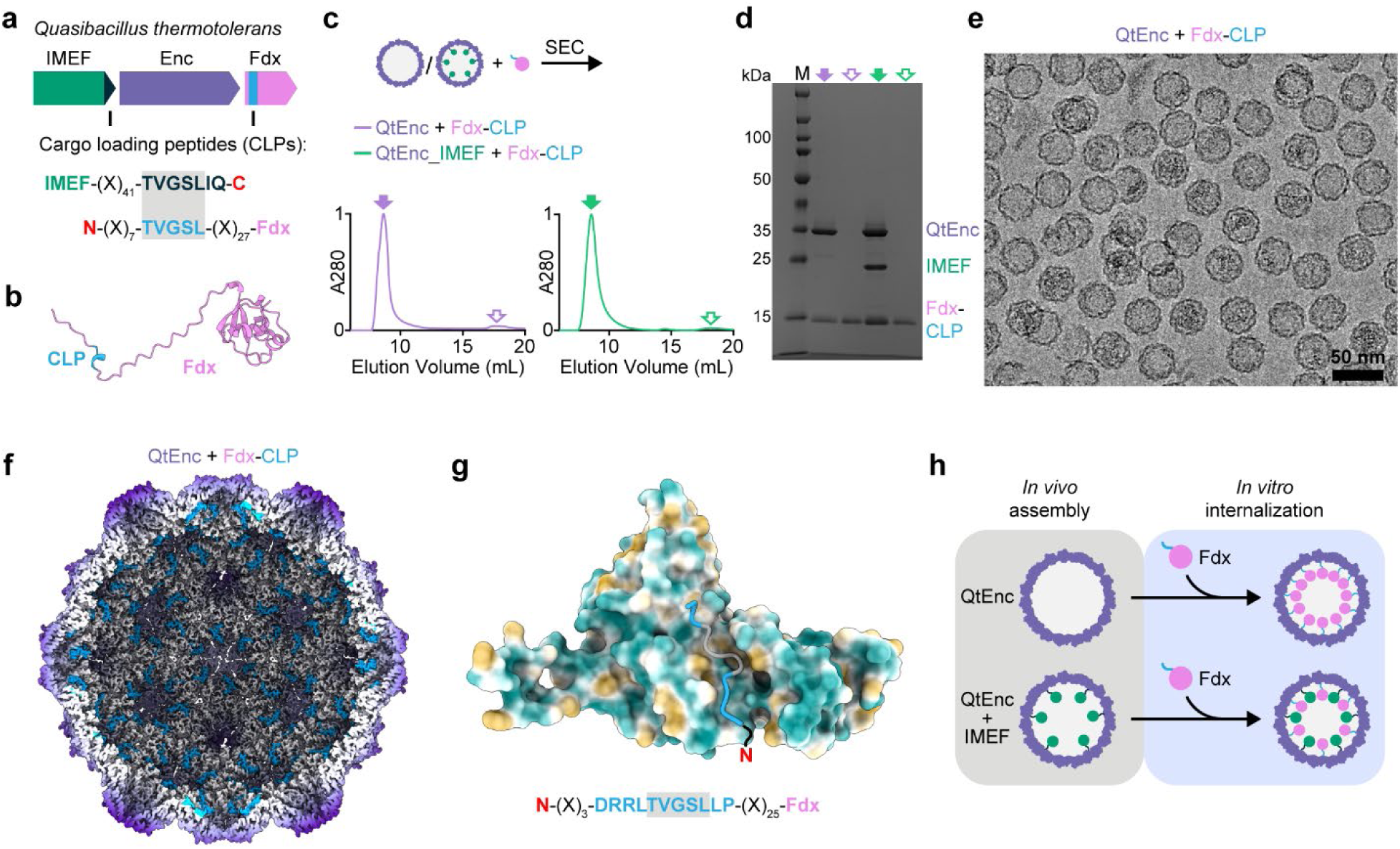
QtEnc shells can internalize proteins *in vitro*. **a**, Schematic of the encapsulin operon in *Quasibacillus thermotolerans*. The operon encodes the cargo IMEF with a C-terminal cargo loading peptide (CLP), the encapsulin shell protein (ENC), and the cargo Fdx with an N-terminal CLP. The core CLP motif (TVGSL) present in both IMEF and Fdx is highlighted. **b**, Alpha Fold predicted structure of Fdx highlighting the CLP located within a long disordered N-terminal extension. **c**, Empty QtEnc and IMEF-loaded QtEnc were incubated with Fdx, followed by size exclusion chromatography (SEC) analysis. Representative normalized SEC traces (purple: QtEnc; green: QtEnc-IMEF) are shown, with filled and outlined arrows indicating the void volume where QtEnc shells elute and free Fdx elution volume, respectively. This experiment was repeated independently three times. A280: absorbance at 280 nm. **d**, Representative SDS-PAGE analysis of fractions corresponding to the indicated arrows in the SEC traces shown in panel c, demonstrating the co-elution of *in vitro* added Fdx with both empty QtEnc and IMEF-loaded QtEnc. This experiment was repeated independently three times. M: protein marker. **e**, Representative cryo-EM micrograph of *in vitro* Fdx-loaded QtEnc highlighting intact and properly assembled shells. **f**, Cutaway view of the cryo-EM map of *in vitro* Fdx-loaded QtEnc, highlighting internal CLP densities in blue confirming Fdx internalization. **g**, Hydrophobic surface representation of a QtEnc protomer with bound Fdx CLP shown in ribbon representation. The core CLP motif (TVGSL) is highlighted in gray and additional resolvable residues in cyan. **h**, Schematic illustrating the post-assembly *in vitro* loading of Fdx into both empty and IMEF-loaded QtEnc shells.

In the course of investigating the function of Fdx within the context of iron release from QtEnc shells, we carried out *in vitro* protein-protein interaction studies using separately purified QtEnc shells and Fdx. *In vitro* incubation of the two proteins (1:1 molar ratio; 30 min at 4°C), followed by size exclusion chromatography (SEC) and SDS-PAGE analysis, indicated that QtEnc and Fdx form a stable complex, as evidenced by the co-elution of Fdx (13.4 kDa) with the QtEnc shell (7.7 MDa) in the SEC void volume (Fig. 1, c and d). Similar results were obtained when testing the interaction between *in vivo* IMEF-loaded QtEnc shells and Fdx, indicating that the apparent QtEnc-Fdx interaction was not disrupted by the presence of the primary QtEnc cargo IMEF. To visualize the QtEnc-Fdx interaction, we carried out single particle cryo-EM analysis of *in vitro* assembled QtEnc-Fdx complexes. Raw micrographs highlighted properly assembled and homogeneous QtEnc shells (Fig. 1e). To our surprise, no indication of externally bound Fdx could be detected in the resulting 2.47 Å cryo-EM map (Fig. 1f and Supplementary Fig. 1). Instead, strong density for the Fdx CLP, located at all conserved interior CLP binding sites within the QtEnc shell, could be observed (Fig. 1g). The presence of internal Fdx CLP density suggests that Fdx has gained access to the interior of the QtEnc shell. Notably, the QtEnc shell contains pores only at its three-fold and five-fold symmetry axes, with the largest pore, at the 3-fold axis, measuring ca. 7.2 Å at its narrowest point.^34^ Because these pores are far smaller than Fdx, internalization cannot occur through the pores themselves. As Fdx is the only native encapsulin cargo known to date to carry an N-terminal CLP—all other known cargos possess C-terminal CLPs—our structural analysis represents the first visualization of an N-terminal CLP-shell interaction. While the CLP core motif (TVGSL) of Fdx interacts in a similar manner to that of the primary QtEnc cargo IMEF, the linker positions, connecting the CLPs to their respective globular cargo domains, are reversed. In addition, more CLP residues could be confidently modeled for the Fdx CLP-shell interaction than for the previously reported IMEF CLP-shell interaction, 11 residues (Fdx) vs 7 residues (IMEF), potentially due to the reversed linker attachment within the Fdx CLP.^34^ Taken together, these results suggest that Fdx can be efficiently internalized into apparently assembled QtEnc shells *in vitro*, representing a previously unknown cargo loading mode for encapsulin systems (Fig. 1h).

## *In vitro* cargo loading is fast and CLP-dependent

To investigate if the observed *in vitro* internalization of Fdx into QtEnc shells is dependent on its globular ferredoxin domain, its disordered CLP-containing N-terminal extension, or both, a series of test constructs were prepared and subjected to *in vitro* loading experiments (Fig. 2a). To test if the ferredoxin domain alone can mediate cargo loading and entrapment within QtEnc shells, the N-terminal extension was deleted yielding ΔN-Fdx. To determine the importance of the N-terminal extension and the CLP, the full-length N-terminal extension of Fdx, or a shortened variant thereof containing a minimal CLP (11 residues resolved by Cryo-EM), were N-terminally fused to Small Ubiquitin-like Modifier (SUMO) to be used as test cargos. In addition, the C-terminal CLP of the primary QtEnc cargo IMEF—which shares an identical core CLP motif (TVGSL) with the Fdx CLP—was fused to the C-terminus of SUMO. All four constructs were separately incubated with QtEnc shells *in vitro* (2:1 (cargo:QtEnc) molar ratio; 30 min at 4°C) and subsequently subjected to SEC and SDS-PAGE analysis. While no cargo internalization could be observed for ΔN-Fdx, all three CLP-containing SUMO constructs exhibited efficient cargo loading (ca. 65% CLP occupancy) under the tested conditions (Fig. 2a). These results demonstrate that the globular ferredoxin domain is not required for cargo internalization, and that a core CLP motif fused to either the N- or C-terminus of a non-native cargo protein is sufficient for *in vitro* loading.

**Fig. 2.**
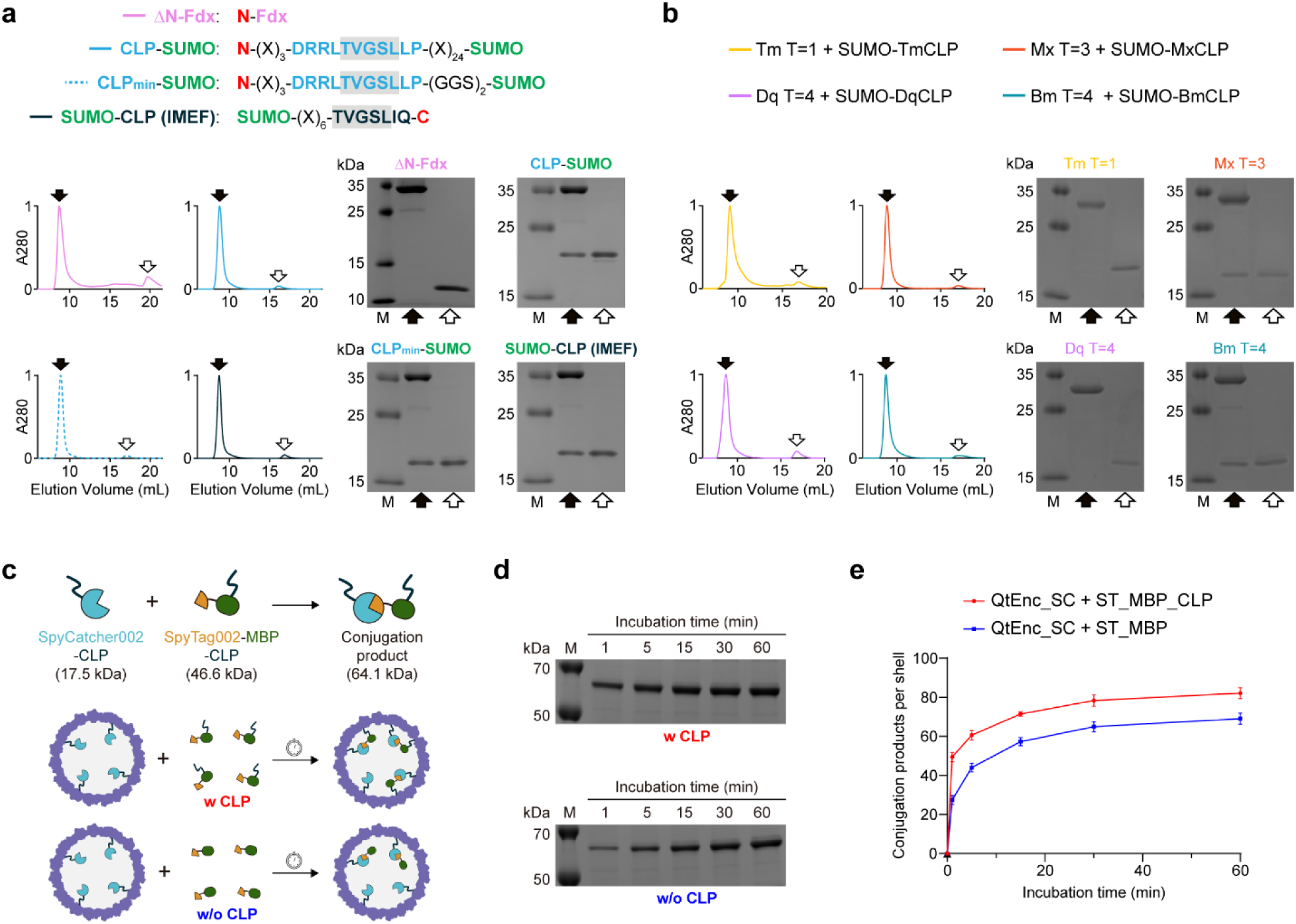
Characterizing determinants of *in vitro* cargo loading. **a**, SEC and SDS-PAGE analyses were used to determine the importance of the presence and position of CLPs in cargos for successful *in vitro* cargo loading. QtEnc was incubated with ΔN-Fdx (N-terminal extension truncated), CLP-SUMO, CLP_min_-SUMO, or SUMO-CLP, followed by SEC analysis (CLP sequences and fusion positions shown on top). Representative normalized SEC traces of loading reactions are shown with filled and outlined arrows indicating the void volume and free cargo elution volume, respectively (bottom left). Representative SDS-PAGE gels of fractions highlighted in SEC traces (bottom right). The results indicate that the core CLP motif is sufficient for *in vitro* cargo loading. All experiments were repeated independently three times. **b**, *In vitro* cargo loading capability was tested across different encapsulin systems, including T=1 (TmEnc) and T=3 (MxEnc) encapsulins with smaller shell sizes, as well as IMEF (BmEnc) and non-IMEF (DqEnc) T=4 encapsulins. SUMO constructs C-terminally fused to their respective native CLPs were used as test cargos. Representative normalized SEC traces are shown, with filled and outlined arrows indicating the void volume and free cargo elution volume, respectively (right). Representative SDS-PAGE gels of the corresponding fractions are shown. The results indicate that MxEnc and BmEnc can be cargo loaded *in vitro*, whereas TmEnc and DqEnc cannot be *in vitro* loaded. All experiments were repeated independently three times. **c**, Schematic illustrating the SpyTag002/SpyCatcher002-based assay used to determine the timescale required for *in vitro* cargo loading into QtEnc. SpyCatcher002-CLP-loaded QtEnc was mixed with SpyTag002-MBP (maltose binding protein), either with or without fused CLP, for defined time periods, followed by SDS-PAGE analysis to quantify formation of the conjugation product. **d**, Representative SDS-PAGE gels showing the formation of the conjugation products over time. Top: loading experiment with cargo carrying a CLP (w CLP). Bottom: loading experiment with cargo not carrying a CLP (w/o CLP). **e**, Quantification of conjugation product formation per shell over time, based on gel densitometry. The results indicate that cargo internalization is not triggered by the presence of an external CLP and that a cargo loading plateau is achieved on a timescale of approximately one hour. Data are shown as mean values, with error bars representing the standard deviation of three independent experiments.

To explore if the ability to internalize cargo *in vitro* is unique to the QtEnc system or potentially also found in other encapsulins, we carried out *in vitro* loading experiments with a selection of diverse structurally characterized encapsulin shells (Fig. 2b). We tested T = 1 (TmEnc),^36^ T = 3 (MxEnc),^37,38^ and other T = 4 (DqEnc and BmEnc)^39^ encapsulins to investigate if shell size is a determinant of *in vitro* loading ability. Further, we assayed if the type of encapsulin system, as determined by the type of native cargo, plays a role in an encapsulin’s *in vitro* loading capability by testing both IMEF (BmEnc) and non-IMEF (DqEnc) T = 4 encapsulin shells.^39^ Using SUMO constructs C-terminally fused to the respective native CLPs of each encapsulin system as test cargos, we found that MxEnc and BmEnc exhibited detectable *in vitro* cargo loading, whereas TmEnc and DqEnc did not (Fig. 2b and Supplementary Fig. 2a). We hypothesize that shell permeability may be linked to the structural dynamics and stability of the shell, and that larger encapsulins with higher triangulation numbers may have a greater propensity to exhibit shell permeability. Beyond this structural consideration, shell permeability may also be related to the native biological function of an encapsulin system. Although QtEnc, BmEnc, and DqEnc are currently the only reported T = 4 encapsulins—making broad generalization difficult—the fact that both T = 4 IMEF encapsulins (QtEnc and BmEnc) exhibit cargo permeability while DqEnc does not, may hint at the possibility that certain encapsulins have evolved distinct shell properties needed to facilitate their native functions, in this case with respect to iron storage or release in T = 4 IMEF encapsulins.

To gain a deeper understanding of the time scale required for *in vitro* loading into QtEnc shells, time course experiments were carried out utilizing a SpyTag/SpyCatcher system—in particular, SpyTag002/SpyCatcher002^40^—to covalently trap internalized cargo (Fig. 2c). In detail, CLP-tagged SpyCatcher was first loaded into QtEnc shells *in vitro* and purified by SEC. Subsequently, SpyTag-maltose binding protein (MBP) fusion constructs, with or without a CLP (SpyTag-MBP-CLP and SpyTag-MBP), were used to probe *in vitro* loading into SpyCatcher-loaded QtEnc shells. Upon internalization, the SpyTag portion of the cargo would be rapidly covalently conjugated to luminal SpyCatcher (SpyTag002/SpyCatcher002 rate constant: 2.0 ± 0.2 × 10^4^ M^−1^ s^−1^),^40^ ensuring that cargo internalization—rather than conjugation—is the rate-limiting step and generating an easily detectable readout in the form of a novel higher molecular weight conjugation product. At defined time points, samples were collected and subjected to SDS-PAGE analysis to quantify conjugation product formation over time (Fig. 2d). A cargo loading plateau was reached within ca. 40 min for both SpyTag-MBP constructs, with the CLP-tagged cargo reaching a higher loading plateau than its untagged counterpart (Fig. 2e and Supplementary Fig. 3). The conjugation observed for the untagged cargo indicates that cargo internalization is not triggered by the presence of an external CLP, confirming that cargo uptake is an intrinsic property of the QtEnc shell.

## Efficient *in vitro* loading can be achieved over a broad cargo size range

To explore the size range of cargos amenable to *in vitro* loading, five proteins spanning a broad range of molecular weights (MW) and sizes—SUMO-CLP (14 kDa, 4 nm), mNeonGreen-CLP (30 kDa, 5.5 nm), Anthrolysin O-CLP (56 kDa, 12 nm), CLP-MerA (mercury reductase A) (126 kDa, 12 nm), and β-galactosidase-CLP (482 kDa, 18 nm)—were used as test cargos (Fig. 3a). All test cargos carried C-terminal IMEF CLP, with the exception of MerA, for which a shortened N-terminal Fdx CLP was used instead because of the catalytic importance of its C-terminus. Respective cargos were incubated with QtEnc shells (2:1 (cargo:QtEnc) molar ratio; 2 h at 4°C), followed by SEC and SDS-PAGE analysis (Fig. 3, b and c). Notably, all tested cargos could be successfully loaded into QtEnc shells, even the very large tetrameric β-galactosidase complex as visualized via negative-stain transmission electron microscopy (Fig. 3d). Dynamic light scattering confirmed that the cargo-loaded shells remained monodisperse and intact across all five cargos (Supplementary Fig. 2b), with loading occupancy being inversely proportional to cargo MW and size (Fig. 3e). The fact that large cargos can be loaded *in vitro* highlights the generalizability of this newly discovered *in vitro* cargo loading approach and also implies that substantial, yet transient structural changes must occur within the QtEnc shell to allow the internalization of such large protein complexes.

**Fig 3.**
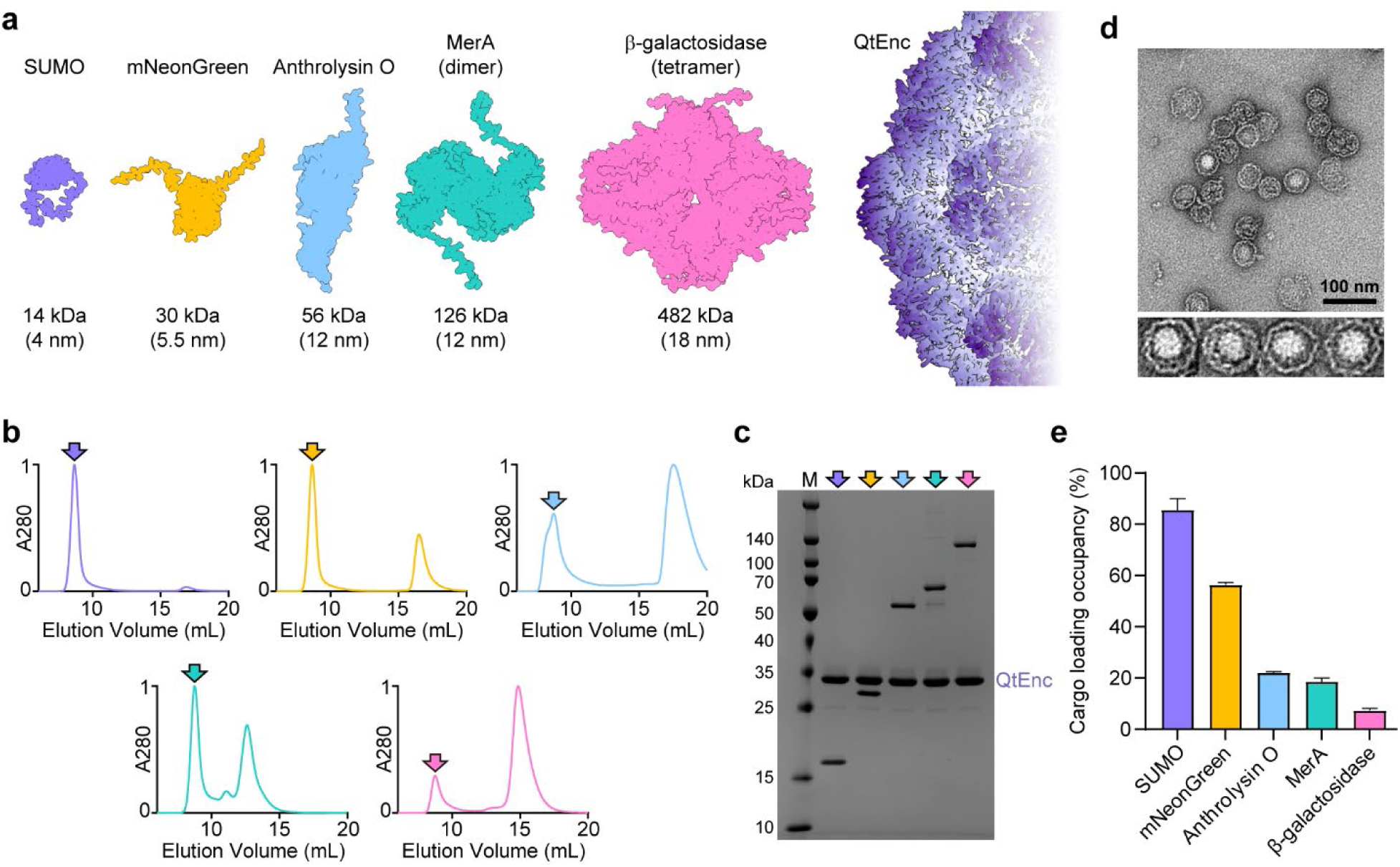
Investigating the influence of cargo size on *in vitro* cargo loading. **a**, Structures of the five cargos tested for *in vitro* loading, ranging from 14 kDa to 482 kDa and 4 to 18 nm in size, shown in surface representation. A section of the QtEnc shell is shown on the right for scale comparison. **b**, Representative normalized SEC traces for *in vitro* loading experiments of the five test cargos. Filled colored arrows, matching the cargo colors shown in panel a indicate the void volume. Experiments were repeated independently three times. **c**, Representative SDS-PAGE gel of void fractions demonstrating that all tested cargos were successfully loaded. This experiment was repeated independently three times. **d**, Representative negative stain micrograph of β-galactosidase-loaded QtEnc highlighting large, internalized protein densities. **e**, Quantification of cargo loading based on CLP binding site occupancy (one binding site per shell protein) of each tested cargo, determined by gel densitometry. The results indicate that cargo loading occupancy is inversely proportional to cargo molecular weight and size. Data are shown as mean values, with error bars representing the standard deviation of three independent experiments.

This raises the question of how the shell accommodates these changes while maintaining overall structural integrity. To investigate, we asked whether QtEnc permeability arises from an assembly–disassembly equilibrium, in which subunits freely dissociate and exchange between shells. Mixing a C-terminally His-tagged QtEnc variant (QtEnc-His) with untagged QtEnc, followed by Ni-NTA affinity chromatography, revealed that the large majority of untagged QtEnc did not associate with QtEnc-His, with only a small fraction co-eluting; this co-eluting fraction increased at higher shell concentrations, where the frequency of shell–shell contacts is greater (Supplementary Fig. 4). These results indicate that subunit exchange between shells is a rare and possibly contact-dependent event rather than a consequence of free, steady-state subunit dissociation.

To directly probe the loading process, we further performed cryo-electron tomography (cryo-ET) on QtEnc shortly after addition of the large β-galactosidase-CLP cargo. The tomograms revealed both empty and β-galactosidase-loaded shells, the vast majority of which appeared intact (Supplementary Fig. 5, a and b, Supplementary Movie 1). We were also able to capture a small number of shells in a partially open state, some engaging cargo-like densities at the site of opening, without any large-scale shell fragmentation or disassembly (Supplementary Fig. 5b). Together, these observations support a model in which QtEnc permeability arises not from a global assembly–disassembly equilibrium, but from local, reversible opening of shell elements that remain attached to the cage, analogous to a hinged lid, transiently admitting cargo before resealing.

## Controlled multiplexed *in vitro* loading of QtEnc shells

We next set out to test if multiple distinct cargos could be simultaneously co-loaded *in vitro* and if their relative loading ratios could be modulated by simply varying their relative amounts in the loading mixture. Utilizing C-terminal CLP-tagged mTagBFP2 and mNeonGreen as test cargos, co-loading experiments were carried out at three different cargo ratios (mTagBFP2:mNeonGreen:QtEnc = 4:2:6, 3:3:6, and 2:4:6) (Fig. 4a). The resulting samples were purified by SEC and analyzed by absorbance-based quantification of fluorescent proteins alongside SDS-PAGE analysis (Fig. 4b). We found that both cargos could be simultaneously internalized into QtEnc shells and that the apparent ratio of co-loaded cargos could be modulated by simply adjusting their proportions in the *in vitro* loading mixture. To confirm that true co-loaded shells are produced—and not two separate populations of mTagBFP2- and mNeonGreen-loaded shells—Förster resonance energy transfer (FRET) experiments were carried out. Co-localization of both cargos within the QtEnc shell lumen would place mTagBFP2 (λ_Ex_ = 399 nm; λ_Em_ = 454 nm) in close proximity to mNeonGreen (λ_Ex_ = 506 nm; λ_Em_ = 517 nm) enabling FRET, with mTagBFP2 acting as the donor and mNeonGreen as the acceptor. Upon excitation at 399 nm and emission detection at 517 nm, a clear FRET signal—ca. four-fold higher than background—was detected in the co-loaded sample (mTagBFP2:mNeonGreen:QtEnc protomer = 3:3:6) but not in control samples (Fig. 4c). To further explore the multiplexing capacity of QtEnc *in vitro* loading, the number of co-loaded cargos was increased to three by adding mCherry (λ_Ex_ = 587 nm; λ_Em_ = 610 nm), and three-color FRET (excitation at 399 nm; emission detection at 610 nm) was employed to probe mixed co-localization within QtEnc shells. A clear FRET signal, ca. three-fold above background, was detected, confirming the simultaneous co-internalization of all three distinct cargos (Fig. 4d).

**Fig 4.**
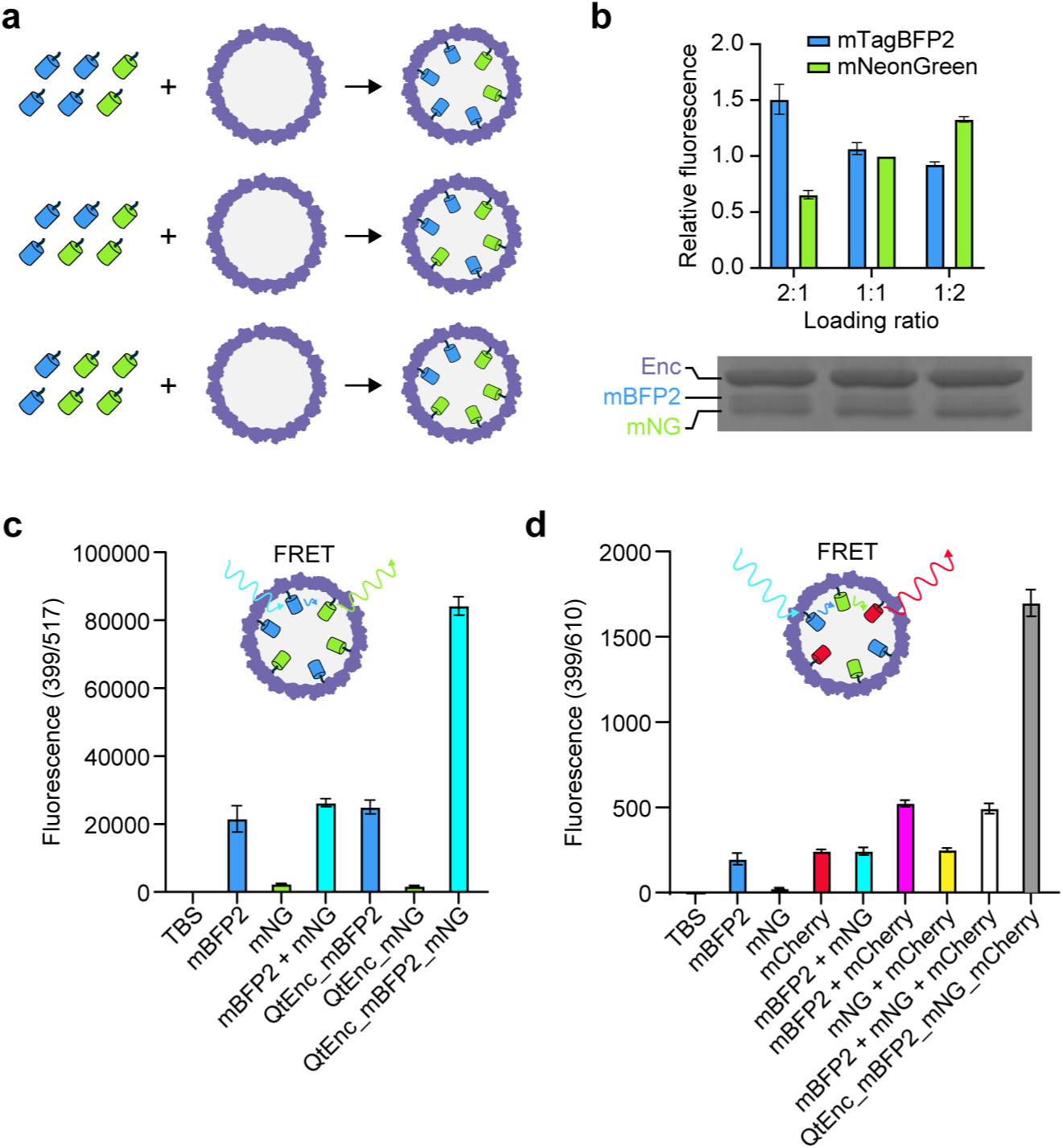
Controlled multiplexed *in vitro* loading of QtEnc shells. **a**, Schematic illustrating the *in vitro* co-loading of mNeonGreen and mTagBFP2 at varying ratios. **b**, Quantification of co-loading reactions showing the relative fluorescence of mNeonGreen and mTagBFP2 for three different co-loading ratio samples, with representative SDS-PAGE analysis of the corresponding samples shown below. The results indicate that the apparent ratio of co-loaded cargos can be modulated by adjusting the input ratios in the *in vitro* loading mixture. Data are shown as mean values, with error bars representing the standard deviation of three independent experiments. **c**, Confirmation of co-loading and co-localization via FRET analysis (excitation: 399 nm, emission: 517 nm) for the indicated samples. Preceding underscores indicate loaded cargos. Data are shown as mean values, with error bars representing the standard deviation of three independent experiments. **d**, Confirmation of triple cargo co-loading and co-localization inside QtEnc. via FRET analysis (excitation: 399 nm, emission: 610 nm) for the indicated samples. Data are shown as mean values, with error bars representing the standard deviation of three independent experiments.

## QtEnc shells stably retain and protect internalized cargo

Because the CLP–shell interaction is non-covalent and the QtEnc shell is permeable, we first asked whether internalized cargo remains stably encapsulated over time or is gradually released. To test this, we loaded His-tagged mNeonGreen into QtEnc and incubated the mNeonGreen-loaded shells for 6, 12, and 24 h in both buffer (TBS) and cell culture medium, after which each sample was applied to a Ni-NTA spin column. In this assay, cargo released from the shell exposes its His-tag and would be captured on the column, whereas encapsulated cargo, with its His-tag occluded by the shell, would not be able to bind the resin. Analysis of the flow-through and elution fractions by SDS-PAGE revealed no detectable cargo release under either condition over the entire time course (Supplementary Fig. 6), indicating that internalized cargo remains stably encapsulated. We attribute this stable retention to the high-affinity CLP–protomer interaction, resulting from shape complementarity between the CLP and an interior surface groove of the protomer together with conserved ionic and hydrophobic interactions.^34^

One of the primary advantages of cargo protein encapsulation is protection from proteolytic degradation and other harsh environmental conditions.^40,41^ This protection is typically attributed to the shell acting as a rigid physical barrier that limits the access of external factors, such as proteases, to the encapsulated cargo. We therefore asked whether the QtEnc shell can still confer protection to encapsulated cargo despite its permeable nature. To this end, free mNeonGreen and mNeonGreen-loaded QtEnc were incubated with the protease trypsin at 37°C, and samples collected at defined time points were analyzed by denaturing gel electrophoresis to assess the extent of mNeonGreen degradation (Supplementary Fig. 7). Within 10 min, free mNeonGreen was completely cleaved, with no intact full-length protein detectable, and was progressively degraded to smaller fragments thereafter. In contrast, full-length mNeonGreen remained detectable even after 2 h of incubation when loaded inside QtEnc. The gradual degradation observed for QtEnc-loaded mNeonGreen further supports the permeability of the QtEnc shell, which allows trypsin to access the encapsulated cargo. We hypothesize that the enhanced protection of QtEnc-loaded mNeonGreen relative to free mNeonGreen results from CLP-mediated tethering of the cargo to the shell interior, which likely substantially limits the protease-accessible, exposed regions of the cargo, rather than from complete physical exclusion of proteases.

## Efficient low pH-triggered cargo release from QtEnc shells

Having characterized the permeable nature of the QtEnc shell, we reasoned that beyond *in vitro* cargo loading, this property would also be useful for efficiently releasing cargo from the QtEnc shell without the need for shell disassembly, if an appropriate way of detaching cargo proteins from their CLPs could be found. Such a system would be particularly useful in the context of nanocarrier design for cytosolic protein delivery. For efficient protein nanocage-based cytosolic protein delivery, an important but often overlooked step is cargo-shell detachment. Without timely detachment, cargos may remain entrapped within endosomes and be trafficked to phagolysosomes, where they are ultimately degraded.^42^ To achieve timely cargo-shell detachment and allow cargo release from the QtEnc shell, we incorporated a point-mutated (G150N) pHIntein (hereafter referred to as pHIntein) into the cargo.^43^ pHIntein is a protein capable of self-cleaving its C-terminus under acidic conditions (pH 5-6). When N-terminally fusing a CLP and C-terminally fusing a protein of interest (POI) to pHIntein, this cleavage would result in the pH-triggered detachment of the POI cargo from the CLP-pHIntein (Fig. 5a). We envisioned that pHIntein-based cargo, with the permeable QtEnc shell, would together allow endosome-specific, acid-triggered release of the POI via pHIntein cleavage, followed by POI escape from the shell facilitated by its intrinsic permeability (Fig. 5b). Specifically, the cargo was designed with the following domain order from the N- to the C-terminus: an N-terminal minimal Fdx CLP, pHIntein, and the POI. Using mNeonGreen as the POI, we first evaluated the acid-triggered cleavage activity of pHIntein. At pH 6 and 37°C, approximately 85% of the unencapsulated cargo was cleaved within 3 h, whereas at pH 7.5 only 28% was cleaved, consistent with the reported pH-triggered cleavage efficiency (Supplementary Fig. 8).^43^ Using the same cargo construct (CLP-pHIntein-mNeonGreen), we then tested whether the permeable QtEnc shell enables not only *in vitro* cargo loading but also POI release. QtEnc loaded with CLP-pHIntein-mNeonGreen was incubated at pH 6 and 37°C for 3 h, followed by SEC to separate released mNeonGreen from the QtEnc shell. The SEC profile, together with the corresponding SDS-PAGE analysis, indicated that the loaded CLP-pHIntein-mNeonGreen was efficiently cleaved, with a cleavage efficiency comparable to that of the free cargo, and that the untethered mNeonGreen was quantitatively released from the shell (Fig. 5c).

**Fig 5.**
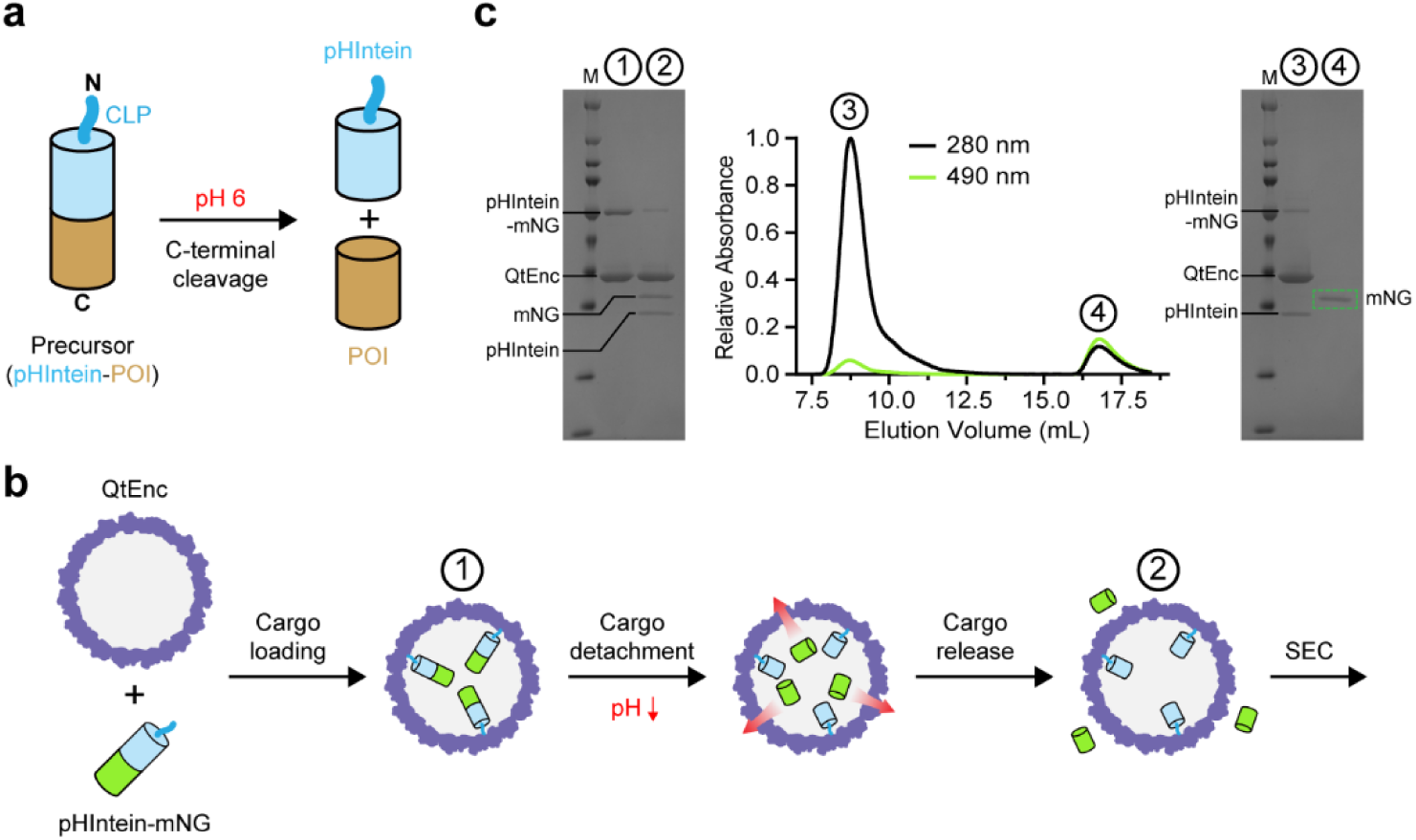
A pHIntein-based cargo detachment module enables low pH-triggered cargo release. **a**, Schematic illustrating the low pH-triggered C-terminal cleavage activity of pHIntein, releasing the C-terminally fused protein of interest (POI). **b**, Schematic illustrating the steps of POI (mNeonGreen: mNG) loading, detachment, and release from the QtEnc shell under acidic conditions, mediated by QtEnc shell permeability and pHIntein cleavage activity. **c**, The numbered steps shown in panel b were examined using SDS-PAGE analysis and SEC. Representative gels and SEC traces are shown. Samples in the SDS-PAGE gels are annotated with numbers (1: cargo loaded QtEnc, 2: sample after low pH incubation and before SEC separation, 3: QtEnc-containing void SEC fraction, 4: released mNG-containing fraction, no longer associated with QtEnc) corresponding to the labeled steps and SEC fractions. Successful *in vitro* cargo loading, cargo detachment, and release from the QtEnc shell were confirmed using pHIntein–mNeonGreen as a proof-of-concept cargo. Experiments were replicated three times independently.

## A rationally designed QtEnc-based nanocarrier for cytosolic protein delivery

Given the downstream goal of cytosolic delivery, we further modified the cargo by incorporating fusogenic peptides to promote endosomal escape of the untethered and released POI. Two different fusogenic peptides, TAT-S19 and GALA3, were evaluated by inserting the fusogenic peptide between pHIntein and mNeonGreen (Fig. 6a).^44,45^ SEC-based POI release experiments were then repeated using QtEnc loaded with cargos containing the respective fusogenic peptides (Fig. 6, b and c). Both cargo constructs exhibited efficient pHIntein cleavage. Whereas untethered GALA3-mNeonGreen was quantitatively released from the shell (Fig. 6c), untethered TAT-S19-mNeonGreen unexpectedly remained trapped within the shell (Fig. 6b). We reason that this outcome arises from the aggregation propensity of the TAT-S19 peptide,^44^ which likely causes the TAT-S19-mNeonGreen to form large aggregates inside the shell, thereby preventing its escape.

**Fig 6.**
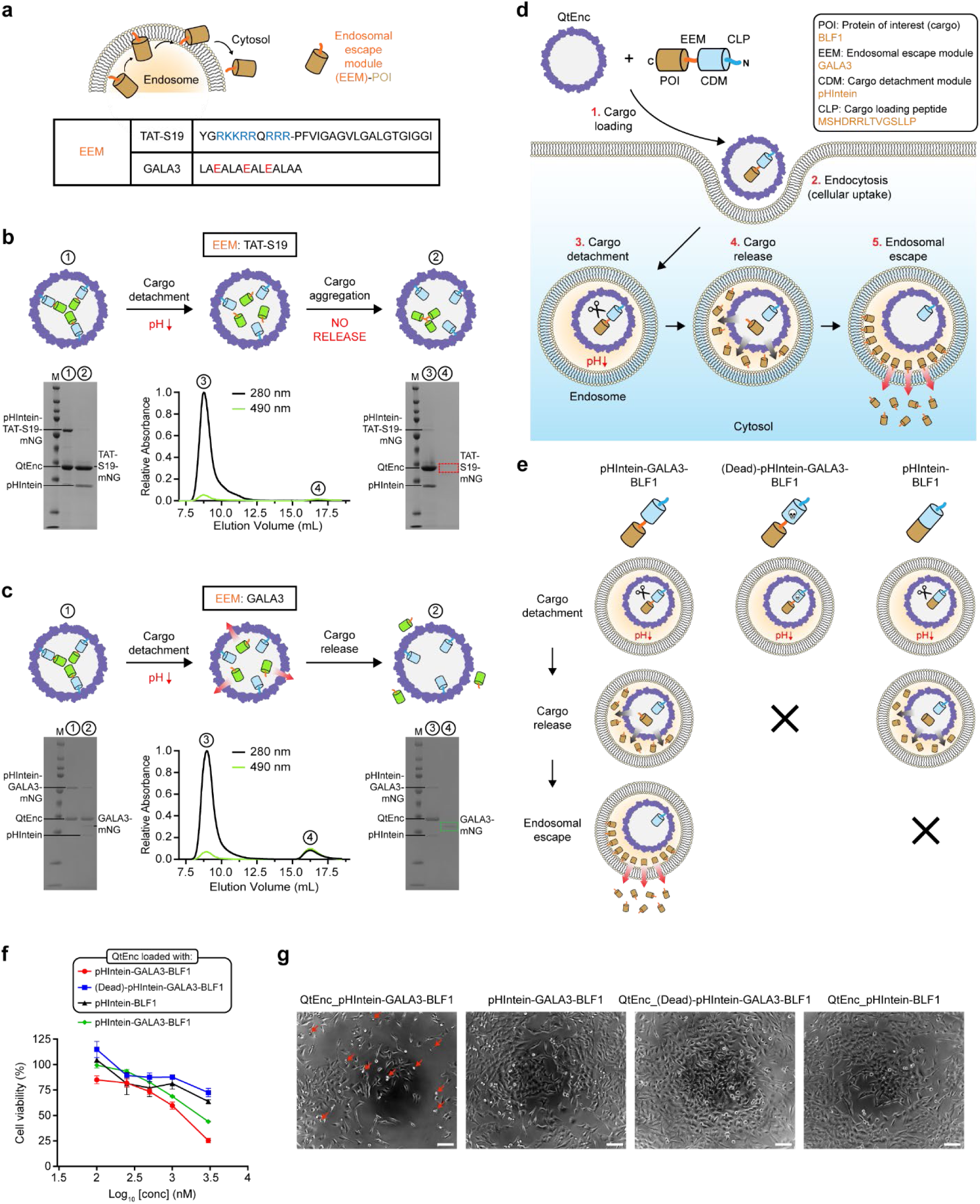
A rationally designed QtEnc-based nanocarrier for cytosolic protein delivery. **a**, Schematic illustrating the role of the endosomal escape module (EEM) in delivering the POI to the cytosol. Sequences of the two EEMs tested in this study, TAT-S19 and GALA3, are shown in the table, with charged residues highlighted. **b**-**c**, Low pH cargo release tests performed using TAT-S19 (b) and GALA3 (c) as EEMs, with mNeonGreen as the protein of interest (POI). Representative SDS-PAGE gels and SEC traces indicate that mNeonGreen was successfully released only when GALA3 was used as an EEM, highlighting the importance of selecting an EEM that does not tend to aggregate under high local molarities or low pH. Experiments were repeated three times independently. **d**, Schematic illustrating the rationally designed QtEnc-based nanocarrier for cytosolic protein delivery and the individual steps involved. **e**, Schematic illustrating the importance of the tested negative controls for the cargo detachment module (CDM, pHIntein) and EEM (GALA3). Inactivation of either module results in failure of cytosolic delivery of the POI (Burkholderia Lethal Factor 1: BLF1). **f**, Cell viability quantified by PrestoBlue-based assays on HeLa cells incubated for 72 h with QtEnc loaded with pHIntein-GALA3-BLF1, (Dead)-pHIntein-GALA3-BLF1, or pHIntein-BLF1, or with free non-encapsulated pHIntein-GALA3-BLF1, at BLF1 concentrations of 0.1, 0.25, 0.5, 1, and 3 µM. QtEnc loaded with the complete pHIntein-GALA3-BLF1 construct elicited the greatest cytotoxic effect across all concentrations, exceeding both the module-deficient controls and the free cargo, confirming that the QtEnc shell, the cargo detachment module (CDM), and the endosomal escape module (EEM) each contribute to efficient cytosolic delivery. Data are shown as mean values, with error bars representing the standard deviation of three independent experiments. **g**, Representative bright-field images of HeLa cells taken at the 72 h time point after incubation with 3 µM (with respect to BLF1) QtEnc loaded with pHIntein-GALA3-BLF1, (Dead)-pHIntein-GALA3-BLF1, or pHIntein-BLF1, or with free non-encapsulated pHIntein-GALA3-BLF1. Red arrows indicate cells displaying morphological changes characteristic of cytotoxicity. Scale bars, 100 µm.

Having demonstrated the utility of the permeable QtEnc shell as a pH-triggered POI release system, we next designed a QtEnc-based NanoCarrier (QtEncNC) as a modular platform for cytosolic protein delivery (Fig. 6d). The QtEncNC test cargo was designed with the following domain architecture, arranged from N- to C-terminus: a CLP for *in vitro* loading into the QtEnc shell (here: the minimal N-terminal Fdx CLP), a Cargo Detachment Module (CDM) for stimulus-triggered POI untethering (here: pHIntein), an Endosomal Escape Module (EEM) to promote endosomal escape of the released POI (here: GALA3), and the POI itself, which can be any therapeutic protein that exerts its effect in the cytosol. As a test POI, we selected the cytotoxic protein Burkholderia Lethal Factor 1 (BLF1) which irreversibly modifies the eukaryotic translation initiation factor eIF4a and needs to reach the cytosol to exert its effect.^46^ The QtEncNC mechanism of action is initiated by endocytosis of cargo-loaded QtEnc shells, consistent with the general propensity of protein nanocages in this size range to be taken up by cells via non-specific endocytic pathways.^10^ Supporting this, fluorescence imaging of HeLa cells incubated with mNeonGreen-loaded QtEnc showed colocalization of mNeonGreen with the acidic-compartment marker LysoTracker, confirming that QtEnc is endocytosed and trafficked to acidic endo/lysosomal compartments (Supplementary Fig. 9). Following endocytosis, endosomal acidification triggers pHIntein self-cleavage, detaching the EEM-fused POI from the shell-bound pHIntein. Subsequently, the EEM-fused POI escapes the QtEnc shell via its inherent permeability, then escapes the endosome and enters the cytosol through the action of the EEM. As a proof of concept, we constructed the QtEncNC test cargo in the following domain arrangement: (N)-CLP-pHIntein-GALA3-BLF1-(C). To validate this design, we first confirmed that GALA3-BLF1 can quantitatively escape the QtEnc shell upon pHIntein cleavage, with no indication of aggregation (Supplementary Fig. 10b). To assess the individual contributions of the CDM and EEM to cytosolic delivery, two control constructs were prepared (Fig. 10e). To evaluate the importance of the CDM, a catalytically inactive pHIntein variant was generated by introducing a point mutation ((Dead)-pHIntein), which abolishes acid-triggered self-cleavage and thereby prevents detachment and shell escape of the EEM-fused POI (Supplementary Fig. 10, a and c). To evaluate the importance of the EEM, GALA3 was omitted from the construct. Although acid-triggered POI detachment and subsequent shell escape are expected to proceed normally (Supplementary Fig. 10d), endosomal escape of the shell-released POI would be less efficient or prevented in the absence of the EEM. Three QtEncNC variants, each loaded with one of the three cargo constructs, were prepared and incubated with HeLa cells at varying concentrations, normalized to the amount of BLF1. After 72 hours, cell morphology was assessed by bright-field microscopy and cell viability was measured via PrestoBlue assay. Cell viability results revealed a clear and concentration-dependent cytotoxic effect for QtEncNC carrying the complete cargo construct (CLP-pHIntein-GALA3-BLF1), whereas both control constructs exhibited attenuated cytotoxicity across all tested concentrations (Fig. 6f). At 1 µM BLF1, the complete construct reduced cell viability to ca. 60%, while the CDM-deficient control ((Dead)pHIntein-GALA3-BLF1) and the EEM-deficient control (pHIntein-BLF1) retained cell viabilities of ca. 88% and ca. 81%, respectively. The difference became even more pronounced at 3 µM, where the complete construct reduced cell viability to ca. 25%, compared to ca. 73% and ca. 64% for the CDM- and EEM-deficient controls, respectively. To assess whether encapsulation itself confers a delivery advantage, we additionally tested the free, non-encapsulated cargo (CLP-pHIntein-GALA3-BLF1) under identical conditions at matched BLF1 concentrations. Across all tested concentrations, the free cargo elicited consistently lower cytotoxicity than QtEnc loaded with the complete construct (Fig. 6f), demonstrating that encapsulation within QtEnc improves the cytosolic delivery efficiency of the cargo. This advantage likely reflects the efficient endocytic uptake of QtEnc shells (Supplementary Fig. 9), whereas the free cargo, lacking a dedicated delivery vehicle, is internalized less efficiently. Bright-field microscopy of cells treated at 3 µM further corroborated these findings, revealing marked morphological changes consistent with cytotoxicity in cells treated with the complete construct, but not in those treated with any control (Fig. 6g). Collectively, these results demonstrate that QtEncNC represents an effective and modular platform for cytosolic protein delivery, in which encapsulation, the CDM, and the EEM each contribute to optimal function.

## Conclusion

In this study, we report that select encapsulin shells, including QtEnc, exhibit unexpected permeability properties that enable fundamentally different modes of cargo loading and release when compared to previously described protein nanocage systems. Existing approaches for loading non-native cargo proteins into protein nanocages have relied on shell disassembly under harsh conditions followed by reassembly,^47^ co-expression strategies requiring careful optimization of expression ratios and/or timing between shell protomer and cargo induction,^20^ or the use of additional triggering components to initiate assembly.^48^ In contrast, the permeable nature of QtEnc enables simple, rapid, and single-step *in vitro* cargo loading by mixing the QtEnc shell with a cargo bearing a short CLP core motif fused to either its N- or C-terminus. The minimal size of the required CLP core motif (5 residues) is a particularly attractive feature, as it is unlikely to interfere with the folding or function of the cargo protein. Furthermore, *in vitro* cargo internalization into QtEnc shells reaches a loading plateau within hours at 4°C. The broad applicability of this *in vitro* loading approach was demonstrated by the successful encapsulation of cargo proteins spanning a wide size range, from 14 kDa to 482 kDa. Additionally, multiplexed cargo co-encapsulation with tunable loading ratios was readily achieved, a feature that could be particularly valuable for applications such as multi-enzyme nanoreactors requiring the co-localization of multiple catalytic components. Despite its permeable nature, QtEnc was also shown to confer substantial proteolytic protection to encapsulated cargo.

Building on these discoveries, we leveraged the permeable nature of QtEnc to develop a modular QtEnc-based NanoCarrier (QtEncNC) for cytosolic protein delivery—a strategy that circumvents the need to engineer nanocages for endosomal stimulus-triggered disassembly, which has remained a major challenge in the field.^47^ By incorporating pHIntein as a cargo detachment module, we demonstrated that the permeability of the QtEnc shell can be exploited to release untethered cargo upon acidification. An important cargo design constraint identified in this study is that the cargo and any fused fusogenic peptides should not exhibit a strong aggregation propensity, as aggregation can prevent efficient cargo release. Using BLF1 as a model cytotoxic cargo, we demonstrated that QtEncNC achieves cytosolic protein delivery in HeLa cells, and that both the cargo detachment and endosomal escape modules are important components of the delivery system to achieve optimal results.

Together, the discoveries reported here establish QtEnc as a uniquely versatile and modular nanocarrier platform for cytosolic protein delivery, with broad potential across diverse biomedical application areas.

## Methods

### Molecular biology and cloning

All constructs used in this study, with the exception of ΔN-Fdx, CLP_min_-SUMO, and CLP-(Dead)-pHIntein-GALA3-BLF1, were ordered from Integrated DNA Technologies (IDT) as *E. coli* codon-optimized gBlocks. Genes encoding ΔN-Fdx, CLP_min_-SUMO, and CLP-(Dead)-pHIntein-GALA3-BLF1 were generated by overhang PCR using Fdx, CLP-SUMO, and CLP-pHIntein-GALA3-BLF1 genes as templates, respectively (Supplementary Tables 1 and 2). All genes were cloned into multiple cloning site 2 (MCS2) of the pETDuet-1 vector using Gibson Assembly. For the QtEnc and IMEF co-expression system, a two-gene operon consisting of the IMEF gene followed by the QtEnc gene was constructed with an intergenic sequence containing an identical ribosome binding site (RBS) derived from MCS2. *E. coli* BL21(DE3) cells were transformed with assembled plasmids via electroporation, and all constructs were sequence-verified by Sanger sequencing (Eurofins Scientific).

### Protein expression and purification

With the exception of pHIntein-containing constructs, all constructs were expressed using ZYM-5052 autoinduction medium supplemented with ampicillin (100 µg/mL). Briefly, 125 mL of fresh autoinduction medium was inoculated 1:1000 from a 5 mL overnight culture and grown at 30°C for ca. 20 h. For pHIntein-containing constructs, expression was carried out in lysogeny broth (LB) supplemented with ampicillin (100 µg/mL). 500 mL of fresh LB medium was inoculated 1:100 from a 5 mL overnight culture, grown at 37°C to an OD_600_ of 0.4-0.5, and induced with isopropyl β-D-thiogalactoside (IPTG) at a final concentration of 0.05 mM. Following induction, cultures were grown at 18°C for ca. 24 h. Cells were harvested by centrifugation (6,000 g, 12 min, 4°C), and the resulting pellets were stored at -80°C until further use.

Cell pellets from encapsulin-expressing cultures were resuspended in Tris buffer (20 mM Tris, 150 mM NaCl, pH 7.5) at 5 mL per gram of wet cell mass. Lysozyme (0.5 mg/mL), Benzonase® nuclease (25 units/mL), and MgCl₂(1.5 mM) were added, and the suspension was incubated on ice for 20 min. Cells were lysed by sonication at 65% amplitude with a pulse cycle of 10 s on/20 s off for a total of 4.5 min (Model 120 Sonic Dismembrator, Fisher Scientific), and the lysate was clarified by centrifugation (10,000 g, 15 min, 4°C). For TmEnc only, the clarified supernatant was subjected to heat treatment at 70°C for 30 min, followed by centrifugation (25,000 g, 25 min, 4°C) to remove aggregated proteins. Ammonium sulfate was added to the supernatant to a final concentration of 20% (15% for TmEnc) and incubated on ice for 40 min, followed by centrifugation (15,000 g, 15 min, 4°C). The resulting supernatant was collected, and ammonium sulfate was added to a final concentration of 40% (75% for TmEnc), incubated on ice for 40 min, and centrifuged (20,000 g, 15 min, 4°C). The resulting pellet was resuspended in 4 mL Tris buffer (pH 7.5), passed through a 0.2 µm syringe filter, and subjected to size exclusion chromatography (SEC) using a Sephacryl S-500 16/60 column equilibrated in Tris buffer (pH 7.5) at a flow rate of 1 mL/min. Encapsulin-containing fractions, as assessed by SDS-PAGE, were pooled, concentrated, and dialyzed into low-salt Tris buffer (20 mM Tris, pH 7.5) using Amicon centrifugal filter units (100 kDa MWCO). The dialyzed sample was loaded onto a HiPrep DEAE FF 16/10 anion exchange column at a flow rate of 3 mL/min to remove nucleic acid contamination. To remove lipid contamination and endotoxins from all encapsulin preparations, a Triton X-114 phase separation method adapted from Aida et al. was employed with minor modifications.^49,50^ Briefly, Triton X-114 was added to pooled encapsulin fractions resulting from ion-exchange chromatograph to a final concentration of 1% (v/v) and mixed with agitation at 4°C for 15 min. Phase separation was induced by incubating the samples at 37°C for 5.5 min, followed by centrifugation (10,000 g, 5 min, 37°C). The encapsulin-containing aqueous phase was carefully recovered, and this process was repeated twice. Residual Triton X-114 was subsequently removed by incubating the encapsulin-containing aqueous phase with Bio-Beads SM-2 resin (Bio-Rad; 5 g per 25 mL) with agitation for 2 h at room temperature, followed by centrifugation (5,000 g, 5 min, room temperature). The resulting supernatant was collected, passed through a 0.2 µm syringe filter, and subjected to SEC using a Superose 6 10/300 GL column equilibrated in Tris buffer (pH 7.5) at a flow rate of 0.5 mL/min. Purified encapsulins were stored in Tris buffer (pH 7.5) at 4°C until further use.

All His-tagged constructs were purified by nickel IMAC followed by SEC in Tris buffer (pH 7.5). With the exception of β-galactosidase-CLP, all His-tagged constructs were subjected to SEC using a Superdex 200 10/300 GL column; a Superose 6 Increase 10/300 GL column was used for β-galactosidase-CLP owing to its larger molecular size. Purified proteins were quantified by A280 (absorbance at 280 nm) using a NanoDrop spectrophotometer and the theoretical extinction coefficient of each respective protein, flash-frozen in aliquots using liquid nitrogen, and stored at -80°C until further use. Full SDS-PAGE gels of all protein purifications are shown in the Supplementary Information (Supplementary Fig. 11).

### SDS polyacrylamide gel electrophoresis

SDS-polyacrylamide gel electrophoresis (SDS-PAGE) was performed using an Invitrogen XCell SureLock Mini-Cell system with Novex 14% Tris-Glycine Mini Protein Gels and SDS running buffer. Samples were mixed with 4X SDS sample buffer, heated at 95°C for 4 min, briefly centrifuged, and loaded onto the gel. Electrophoresis was carried out at a constant voltage of 225 V for 42 min at room temperature. The Spectra Multicolor Broad Range Protein Ladder (Thermo Fisher Scientific) was used as a molecular weight marker.

### Dynamic light scattering (DLS) analysis

All hydrodynamic size and polydispersity measurements were performed using an Uncle instrument (Unchained Labs) at 15°C in triplicate. Prior to analysis, all encapsulin samples were diluted to 0.4 mg/mL in Tris buffer (pH 7.5) and clarified by centrifugation (10,000 g, 10 min, 4°C).

### Negative-stain transmission electron microscopy (TEM)

Encapsulin samples were diluted to 0.15 mg/mL in Tris buffer (pH 7.5) for negative-stain TEM analysis. Gold grids (200-mesh, Formvar-carbon coated, EMS #FCF200-Au-EC) were rendered hydrophilic by glow discharge at 5 mA for 60 s using an easiGlow system (PELCO). A 4 µL aliquot of the sample was applied to the grid and incubated for 1 min, blotted with filter paper, and briefly washed with 0.75% uranyl formate solution. Grids were then stained with 0.75% uranyl formate for 1 min, blotted with filter paper, and allowed to dry for at least 20 min prior to imaging. TEM micrographs were acquired using a Tecnai T12 electron microscope at the University of Michigan Life Sciences Institute.

### Single particle cryo-electron Microscopy (cryo-EM)

#### Sample preparation

A purified sample of *in vitro* Fdx-loaded QtEnc was concentrated to 3.5 mg/mL in 150 mM NaCl, 25 mM Tris pH 7.5. 3.5 µL of protein sample were applied to freshly glow discharged Quantifoil R1.2/1.3 Cu 200 mesh grids and prepared by plunge freezing in liquid ethane using an FEI Vitrobot Mark IV (100% humidity, 22°C, blot force 20, blot time 4 seconds, wait time 0 s). The grids were immediately clipped and stored in liquid nitrogen until data collection.

#### Data collection

Cryo-EM movies were collected using a ThermoFisher Scientific Titan Krios G4i cryo-electron microscope operating at 300 kV equipped with a Gatan K3 direct electron detector with a BioQuantum imaging filter. 1,544 movies were collected from a single grid using the SerialEM^51^ software package at a magnification of 105,000x, pixel size of 0.834 Å, defocus range of -1.0 µm to -1.8 µm, exposure time of 2.32 s, frame time of 38 ms, and a total dose of 57.2 e^-^/Å^2^.

#### Data processing

CryoSPARC 4.6.2^52^ was used to process the dataset (Supplementary Fig. 1). 1,544 movies were imported, motion corrected by patch motion correction, and the CTF fit was estimated using patch CTF estimation. Exposures with CTF fit resolutions worse than 6 Å were discarded from the dataset, resulting in 1,433 remaining movies. 199 particles were selected manually and used to create templates for template-based particle picking. Template picker was then used to select 78,442 particles, which were then extracted using a box size of 630 pixels. The particles were then downsampled to a box size of 480 pixels and subsequently sorted by two rounds of 2D classification, resulting in 70,828 remaining particles. An initial volume was created by abinitio reconstruction using three classes and I symmetry, resulting in a majority class containing 70,540 particles. These particles were then used for homogeneous refinement against the ab-initio map with I symmetry imposed, per-particle defocus optimization, per-group CTF parameterization, and Ewald sphere correction enabled using a positive curvature sign, resulting in a 2.47 Å map.

#### Model building

For building the model of *in vitro* Fdx-loaded QtEnc, a starting model containing a single protomer of QtEnc and QtFdx in complex was generated using AlphaFold3.^53^ The QtFdx model was truncated to only include the targeting peptide and the QtEnc-Fdx-CLP complex was manually placed into the map using ChimeraX v.1.8,^54^ followed by improved map fit using the fit-in-map command. This step was repeated for three additional QtEnc-Fdx-CLP models, resulting in a complete asymmetric unit containing four models of QtEnc-Fdx-CLP. The model was then manually refined against the density map using Coot v0.9.8.1.^55^ Phenix v 1.20.1-4487-000^56,57^ was then used to further refine the model by real-space refinement with three macrocycles, minimization_global enabled, local_grid_search enabled, and adp refinement enabled. NCS operators were then identified from the map using map_symmetry and applied to the model using apply_ncs to generate the icosahedral shell. The NCS-expanded shell was then refined again using real-space refinement with three macrocycles, minimization_global enabled, local_grid_search enabled, adp refinement enabled, and NCS constraints enabled. The BIOMT operators were identified using the find_ncs command and manually placed into the header of the .pdb file containing a single ASU of the NCS-refined model (Supplementary Table 3).

### Cryo-electron tomography (cryo-ET)

Sample preparation: Purified QtEnc and β-galactosidase-CLP were mixed at a QtEnc protomer:β-galactosidase-CLP molar ratio of 10:1 (final QtEnc concentration 1 mg/mL) in 150 mM NaCl, 25 mM Tris pH 7.5 and incubated for 15 min prior to plunge freezing. 3.5 µL of the sample was applied to a freshly glow-discharged Quantifoil R1.2/1.3 Cu 200 mesh grid and plunge-frozen in liquid ethane using an FEI Vitrobot Mark IV (100% humidity, 22 °C, blot force 20, blot time 4 s, wait time 0 s). Grids were clipped and stored in liquid nitrogen until data collection.

#### Data collection

Tilt series were collected on a Titan Krios cryo-electron microscope operating at 300 kV, equipped with an energy filter (slit width 20 eV), using SerialEM. Tilt series were acquired using a dose-symmetric scheme over a tilt range of −60° to +60° with a 2° increment, at a nominal magnification of 42,000× (pixel size 2.12 Å) and a target defocus of −3 µm. Each tilt image was recorded as a 12-frame movie at a dose of ca. 1.98 e⁻/Å² per tilt, for a total dose of ca. 121 e⁻/Å² across the tilt series.

#### Data processing

Movie frames for each tilt image were motion-corrected using alignframes in IMOD. Tilt series were then aligned by patch tracking (no fiducial markers were used) and reconstructed into tomograms by weighted back-projection using the IMOD software package.

### *In vitro* cargo loading assays

*In vitro* cargo loading was performed in Tris buffer (pH 7.5) by mixing QtEnc, or other encapsulin shells (TmEnc, MxEnc, BmEnc, and DqEnc) where indicated, with cargo at a molar ratio of cargo:encapsulin = 2:1 and incubating at 4°C for 2 h unless otherwise stated. The total mixture volume was 500 µL, with a final encapsulin concentration of 0.5-2.5 mg/mL. Following incubation, the mixtures were subjected to SEC using a Superdex 200 10/300 GL column, with the exception of β-galactosidase-CLP-loaded samples, for which a Superose 6 Increase 10/300 GL column was used. For pHIntein-based cargo-loaded QtEnc samples intended for cell-based experiments, the SEC column was pre-equilibrated with phosphate-buffered saline (PBS) to exchange the buffer prior to cell culture use. *In vitro* cargo-loaded encapsulin shells eluted at the void volume of the respective column.

### SpyCatcher-SpyTag conjugation assays

To determine the kinetics of *in vitro* cargo internalization into QtEnc shells, a SpyCatcher-SpyTag conjugation assay was carried out using three reaction setups: (1) SpyCatcher002-CLP-loaded QtEnc with SpyTag002-MBP-CLP, (2) SpyCatcher002-CLP-loaded QtEnc with SpyTag002-MBP, and (3) SpyCatcher002-CLP with SpyTag002-MBP-CLP as a positive control. The concentration of QtEnc and SpyCatcher002-CLP in SpyCatcher002-CLP-loaded QtEnc sample was determined by gel densitometry using a dilution series of QtEnc and SpyCatcher002-CLP at known concentrations run alongside the sample on SDS-PAGE. All reaction mixtures (total volume: 20 µL) were prepared in Tris buffer (pH 7.5) with a final concentration of 5 µM for each component (SpyCatcher002-CLP, SpyTag002-MBP-CLP, or SpyTag002-MBP) and incubated at room temperature. At defined time points (1, 5, 15, 30, and 60 min), aliquots were withdrawn from each reaction mixture and immediately mixed with 4X SDS sample buffer and heated at 95°C to quench the reaction, followed by SDS-PAGE analysis. All reactions were performed in triplicate.

Conjugation product formation was quantified by gel densitometry. The conjugation product band from the SpyCatcher-CLP + SpyTag-MBP-CLP positive control reaction at 60 min was used as a reference standard representing 100% conjugation (Supplementary Fig. 3). The amount of conjugation product formed at each time point in each reaction mixture was calculated relative to this standard using densitometric analysis. The amount of conjugation product formed per shell over time was subsequently calculated by dividing the molar amount of conjugation product by the molar amount of QtEnc shells in each reaction.

### Cargo loading occupancy determination

Cargo loading occupancy for QtEnc loaded with cargos of different size was determined by gel densitometry. A dilution series of mNeonGreen at known concentrations was run on SDS-PAGE to generate a standard curve correlating band intensity to protein mass. Cargo loading occupancy was subsequently calculated as the molar ratio of loaded cargo to QtEnc protomer, with molar concentrations of each derived from the standard curve using their respective band intensities and theoretical molecular weights.

### Ni-NTA-based subunit-exchange assay

To assess whether QtEnc undergoes subunit exchange between shells, a C-terminally His-tagged QtEnc variant (QtEnc-His) was mixed with untagged QtEnc at equimolar amounts and incubated for 30 min at 4°C in Tris buffer (pH 7.5; total volume 500 µL). Two conditions were tested: a standard-concentration reaction (QtEnc and QtEnc-His each at 0.079 mg/mL) and a higher-concentration reaction (QtEnc and QtEnc-His each at 4.15 mg/mL). QtEnc alone and QtEnc-His alone were prepared in parallel as controls. Each mixture was applied to a Ni-NTA spin column (QIAGEN), which was then washed twice with 600 µL of wash buffer (Tris buffer, 25 mM imidazole, pH 8.0) and eluted with 150 µL of elution buffer (Tris buffer, 500 mM imidazole, pH 8.0). The flow-through, wash, and elution fractions were analyzed by SDS-PAGE.

### Förster Resonance Energy Transfer (FRET) analysis

FRET experiments were performed using a Synergy H1 plate reader (BioTek) in black flat-bottom 384-well plates at room temperature, with all samples measured in triplicate. The following samples were prepared for the mTagBFP2 (mBFP2)-mNeonGreen (mNG) FRET experiment: mBFP2 and mNG co-loaded QtEnc, mBFP2-only loaded QtEnc, mNG-only loaded QtEnc, mBFP2, and mNG. The amounts of mBFP2 and mNG in each sample were normalized and confirmed by SDS-PAGE prior to measurement. Each well contained 50 µL of sample in Tris buffer (pH 7.5), with Tris buffer (pH 7.5) alone serving as background. FRET was assessed by exciting samples at 399 nm and measuring emission at 517 nm, corresponding to mNG acceptor emission.

For the mBFP2-mNG-mCherry FRET experiments, an identical setup was employed with mBFP2, mNG, and mCherry co-loaded QtEnc and the corresponding control samples. FRET was assessed by exciting samples at 399 nm and measuring emission at 610 nm, corresponding to mCherry acceptor emission.

### Ni-NTA-based cargo retention assay

To assess whether encapsulated cargo is stably retained or gradually released over time, QtEnc was loaded with His-tagged mNeonGreen as described in the *in vitro* cargo loading section and purified by size-exclusion chromatography. mNeonGreen-loaded QtEnc was incubated at 4°C for 6, 12, and 24 h in either Tris buffer (pH 7.5) or supplemented cell culture medium (DMEM + 10% FBS + 1% PS). At each time point, samples were applied to a Ni-NTA spin column (QIAGEN), washed twice with 600 µL of wash buffer (Tris buffer, 25 mM imidazole, pH 8.0), and eluted with 150 µL of elution buffer (Tris buffer, 500 mM imidazole, pH 8.0). The flow-through, wash, and elution fractions were analyzed by SDS-PAGE. Cargo released from the shell exposes its His-tag and is captured on the column (recovered in the elution fraction), whereas encapsulated cargo, with its His-tag occluded by the shell, appears in the flow-through.

### Low pH-triggered pHIntein cleavage assays

pHIntein-based cargo constructs were mixed with citrate-phosphate buffer (pH 6.0) at a 1:10 (v/v) ratio and incubated at 37°C. At defined time points (1, 2, and 3 h), aliquots were withdrawn from the reaction mixture, immediately mixed with 4X SDS sample buffer, and heated at 95°C prior to SDS-PAGE analysis.

### Low pH-triggered POI release from QtEnc shells

pHIntein-based cargo-loaded QtEnc samples were mixed with citrate-phosphate buffer (pH 6.0) at a 1:10 (v/v) ratio and incubated at 37°C for 3 h. Prior to SEC injection, an aliquot was withdrawn from the mixture, immediately mixed with 4X SDS sample buffer, and heated at 95°C to serve as a pre-SEC input control. The remainder of the mixture was then subjected to SEC using a Superdex 200 10/300 GL column equilibrated in Tris buffer (pH 7.5). The void volume fraction, containing QtEnc shells, and the later eluting fractions, containing released POI (if any), were collected and analyzed together by SDS-PAGE.

### Cell culture experiments

HeLa cells were cultured in Dulbecco’s Modified Eagle Medium (DMEM) supplemented with 10% fetal bovine serum (FBS) and 1% penicillin-streptomycin (PS) at 37°C in a humidified atmosphere containing 5% CO_2_. For cell viability experiments, thawed HeLa cells were allowed to recover for 24 h, washed with Hanks’ Balanced Salt Solution (HBSS), and cultured for an additional 24 h. Cells were then detached using trypsin, counted, and seeded at a density of 2,000 cells per well in 96-well plates (200 µL per well) and allowed to adhere for 24 h prior to sample addition.

### Live-cell fluorescence imaging of QtEnc uptake and trafficking

HeLa cells, cultured as described above, were seeded at 10,000 cells per well in 96-well plates and allowed to adhere for 24 h prior to imaging. mNeonGreen-loaded QtEnc was diluted in supplemented DMEM to a final mNeonGreen concentration of 300 nM (determined by absorbance at 506 nm using an extinction coefficient of 116,000 M^-1^ cm^-1^) and added to the cells, which were incubated for 6 h at 37°C with 5% CO₂. Cells were then washed twice with PBS, stained with Hoechst 33342, and subsequently stained with LysoTracker Red DND-99 (Thermo Fisher Scientific). After a final PBS wash, cells were imaged live on an EVOS M5000 imaging system (Thermo Fisher Scientific).

## Cell viability assays

The following samples were tested in triplicate: DMEM (supplemented with 10% FBS and 1% PS) alone, 0.1% (v/v) Triton X-100 in supplemented DMEM, QtEnc alone, and QtEncNC loaded with one of three cargo constructs: CLP-pHIntein-GALA3-BLF1, CLP-(Dead)-pHIntein-GALA3-BLF1, and CLP-pHIntein-BLF1. The BLF1 concentration in each QtEncNC stock was determined by gel densitometry using a dilution series of the respective BLF1-containing cargo construct at known concentrations run alongside the QtEncNC sample on SDS-PAGE. Each QtEncNC sample was then diluted in supplemented DMEM to a final BLF1 concentration of 100 nM, 250 nM, 500 nM, 1 µM, and 3 µM, and 100 µL of each dilution was added to per well. For the media-only and Triton X-100 controls, 100 µL of supplemented DMEM or 0.1% (v/v) Triton X-100 in supplemented DMEM was added per well, respectively. For the QtEnc-only control, QtEnc in PBS was diluted in supplemented DMEM to a total volume of 100 µL, with the amount of QtEnc matched to that present in the highest concentration QtEncNC sample (3 µM BLF1 equivalent). Following sample addition, cells were incubated at 37°C with 5% CO₂ for 72 h prior to further analysis.

Prior to cell viability assays, bright-field microscopy images of the cells were acquired using an EVOS M5000 Microscopy System (Thermo Fisher Scientific) to assess cell morphology. Cell viability was assessed using the PrestoBlue Cell Viability Reagent (Thermo Fisher Scientific). Following the 72 h incubation, the culture medium was removed, and cells were washed twice with PBS to remove residual sample. PrestoBlue reagent was diluted 1:9 (v/v) in phenol red-free supplemented DMEM and 100 µL was added to each well. The plate was incubated at 37°C with 5% CO₂ for 45 min, then wrapped in aluminum foil and equilibrated at room temperature for 4 h to stabilize the fluorescence signal and protect the reagent from light exposure. Fluorescence was measured at an excitation wavelength of 560 nm and emission wavelength of 590 nm using a Synergy H1 plate reader (BioTek). Cell viability was expressed as a percentage relative to the QtEnc-only control, after subtracting the background fluorescence obtained from Triton X-100-treated positive control wells. No meaningful difference in cell viability was observed between the QtEnc-only and media-only controls.

## Data availability

The cryo-EM maps and structural models of *in vitro* Fdx-loaded QtEnc have been deposited and are publicly available in the Electron Microscopy Data Bank (EMDB-76203) and Protein Data Bank (PDB ID: 11YX). A tilt series movie of the β-galactosidase *in vitro* loading reaction is provided as Supplementary Movie 1.

## Acknowledgements

T.W.G acknowledges funding from NIH (R35GM133325) and NSF (2342136). S.K. acknowledges funding from Asan Foundation (Biomedical Science Scholarship) and University of Michigan Rackham Graduate School (Rackham Predoctoral Fellowship). Research reported in this work was supported by the University of Michigan Cryo-EM Facility (U-M Cryo-EM). U-M Cryo-EM is grateful for support from the U-M Life Sciences Institute and the U-M Biosciences Initiative. We thank Vinson Lam of the U-M Cryo-EM core for help in cryo-electron tomography. Molecular graphics and analyses were performed using UCSF ChimeraX developed by the Resource for Biocomputing, Visualization, and Informatics at the University of California, San Francisco, with support from the National Institutes of Health R01GM129325 and the Office of Cyber Infrastructure and Computational Biology, National Institute of Allergy and Infectious Diseases.

## Author contributions

S.K. and T.W.G. designed the project. S.K., M.P.A. and T.W.G processed and analyzed cryo-EM data. M.P.A. built the structural models. J.A.J. conducted initial protein-protein interaction experiments. S.K. carried out all other laboratory experiments. S.K. and T.W.G. wrote the manuscript. T.W.G. oversaw the project in its entirety.

## Competing interests

The authors declare the following competing financial interests: S.K. and T.W.G. have filed a patent application related to this work.

## Notes

### Summary of Updates

Multiple additional experiments added including: -Ni-NTA-based dynamic subunit-exchange: this experiment directly addresses the question of weather dynamic subunit-exchange plays a major role in the observed QtEnc permeability. Based on our results, we conclude that it does not represent a major contributor to the observed permeability properties. -Cryo-electron tomography of a QtEnc in vitro loading reaction: this experiment captures the loading reaction in as close to native-like conditions as possible. Based on our observations, we formulate a QtEnc permeability model not based on large-scale dynamic shell disassembly/reassembly or subunit exchange, but rather on dynamic shell elements, individual protomers or pentameric/hexameric facets, transiently opening while remaining attached to the QtEnc shell, analogous to a hinged lid, briefly admitting cargo association before resealing. -Ni-NTA-based cargo retention assay: this experiment addresses the question of cargo retention over time. Our results lead us to conclude that in both buffer and cell culture media, over a 24 h period, no detectable cargo release is taking place. -Intracellular trafficking of QtEnc-delivered cargo: this experiment shows that mNeonGreen-loaded QtEnc is quickly internalized into HeLa cells and co-localizes with endolysosomes based on LysoTracker staining, thereby confirming the proposed endocytic trafficking mechanism of the QtEncNC nanocarrier system.

