## Supplementary Information for "A permeable protein nanocage enables facile cargo loading and cytosolic protein delivery"

**Table of Contents**


| | | |
|---|---|---|
| Supplementary Fig. 1 | Cryo-EM data processing workflow for in vitro Fdx-loaded QtEnc | 2 |
| Supplementary Fig. 2 | Dynamic light scattering (DLS) analysis of encapsulin shells and cargo-loaded QtEnc variants | 3-4 |
| Supplementary Fig. 3 | SpyCatcher002-SpyTag002 conjugation positive control | 5 |
| Supplementary Fig. 4 | Ni-NTA-based dynamic subunit-exchange assay of QtEnc and QtEnc-His | 6 |
| Supplementary Fig. 5 | Cryo-electron tomography of QtEnc mixed with β-galactosidase-CLP | 7 |
| Supplementary Fig. 6 | Ni-NTA-based cargo retention assay of mNeonGreen-loaded QtEnc over time | 8 |
| Supplementary Fig. 7 | Protease protection of encapsulated mNeonGreen by the QtEnc shell | 9 |
| Supplementary Fig. 8 | pH-dependent cleavage activity of free pHIntein | 10 |
| Supplementary Fig. 9 | Intracellular trafficking of mNeonGreen-loaded QtEnc in HeLa cells | 11 |
| Supplementary Fig. 10 | Characterization of pHIntein-mediated cargo detachment and shell release for QtEncNC cargo constructs | 12 |
| Supplementary Fig. 11 | Uncropped and annotated SDS-PAGE gels | 13-16 |
| Supplementary Table 1 | Aimo acid sequences of protein constructs used | 17-21 |
| Supplementary Table 2. | DNA primers used in this study | 22 |
| Supplementary Table 3. | Cryo-EM data collection and refinement statistics | 23 |

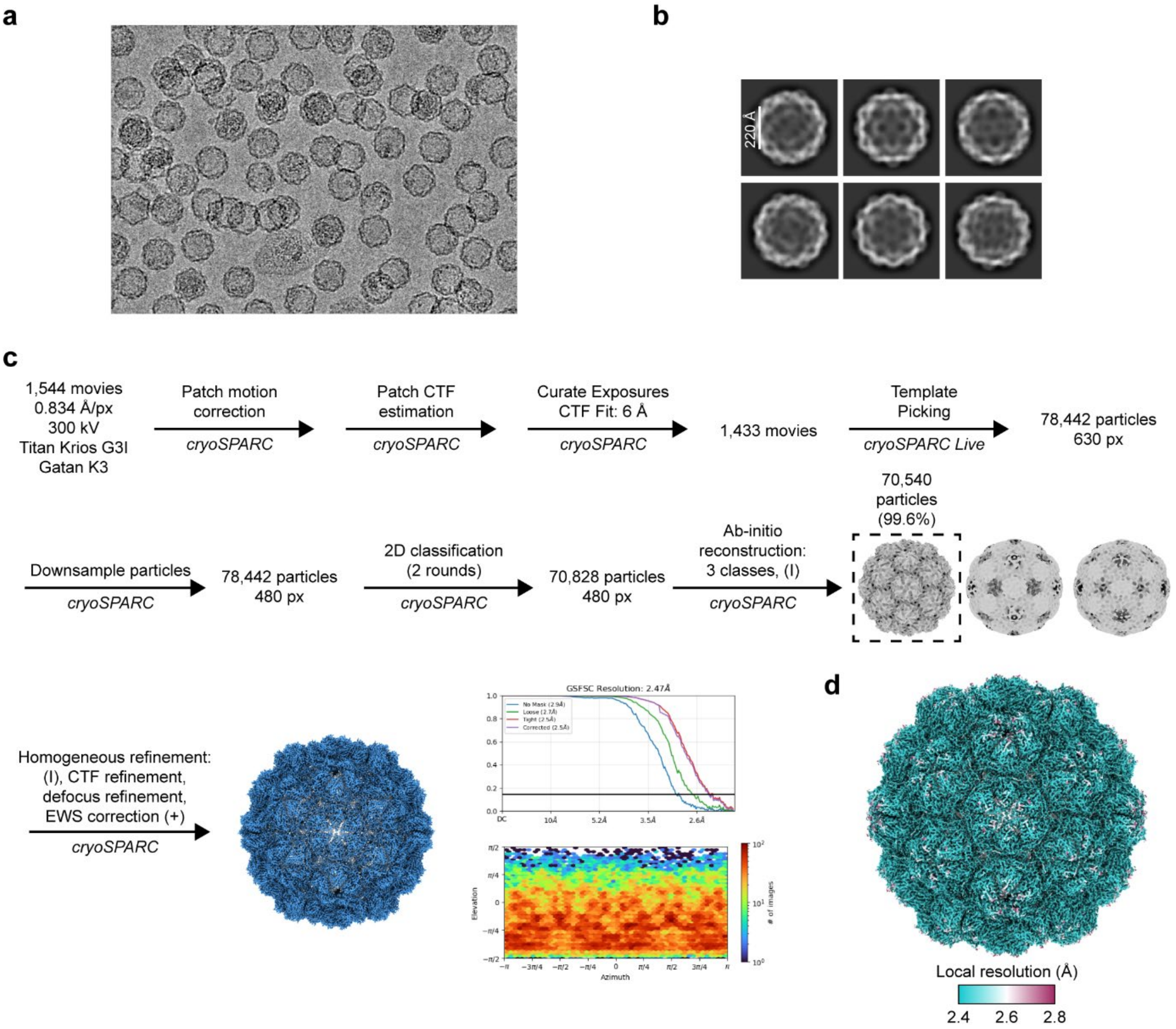


**Supplementary Fig. 1. Cryo-EM data processing workflow for *in vitro* Fdx-loaded QtEnc.** **a**, Representative raw cryo-EM micrograph. **b**, Representative 2D class averages. **c**, Data processing workflow including Fourier Shell Correlation (FSC) plot and viewing direction plot. **d**, Local resolution analysis.

**a**

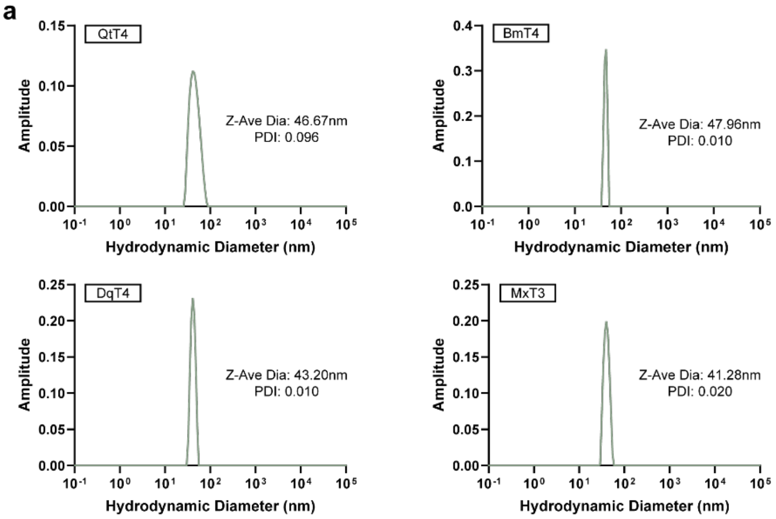


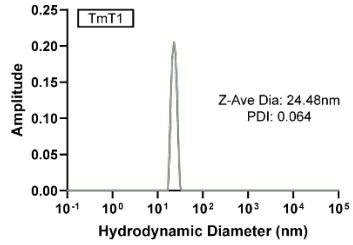


| Triangulation number | Encapsulin | *In vitro* cargo internalization (y/n) |
|---|---|---|
| T=4 | QtEnc | Y |
| | BmEnc | Y |
| | DqEnc | N |
| T=3 | MxEnc | Y |
| T=1 | TmEnc | N |

**b**

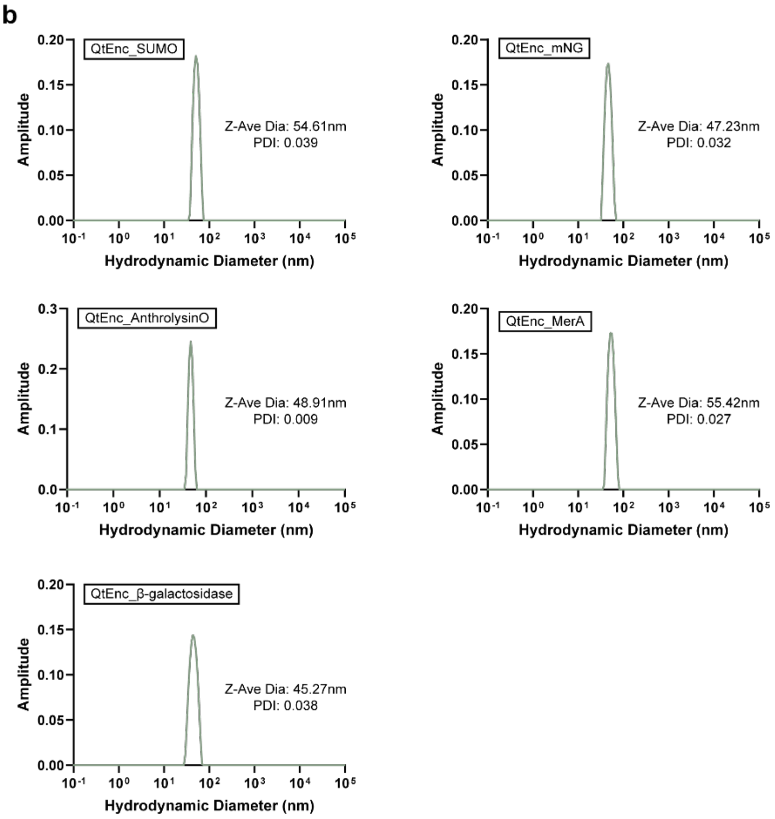


**Supplementary Fig. 2. Dynamic light scattering (DLS) analysis of encapsulin shells and cargo-loaded QtEnc variants. a**, Empty shells: QtEnc, BmEnc, DqEnc, MxEnc, TmEnc. The accompanying table summarizes each encapsulin by triangulation number and its *in vitro* cargo internalization outcome (Y, internalization observed; N, no internalization) **b**, Cargo-loaded shells: QtEnc_SUMO, QtEnc_mNG, QtEnc_Anthrolysin O, QtEnc_MerA, QtEnc_β-galactosidase. The Z-average hydrodynamic diameter and polydispersity index (PDI) are indicated within each plot. All samples exhibit a single, narrow peak with PDI values below 0.1, indicating high monodispersity. No meaningful change in hydrodynamic diameter was observed upon cargo loading relative to empty QtEnc shells.

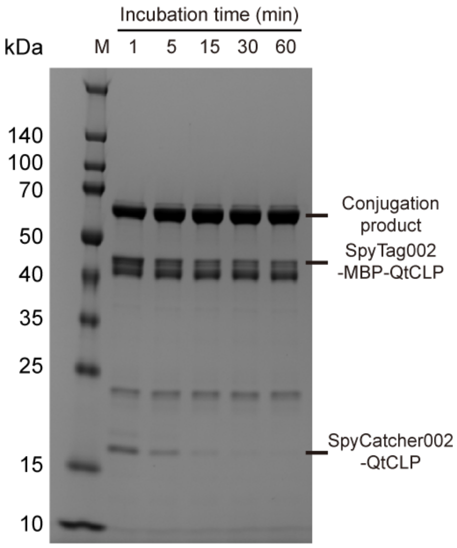


**Supplementary Fig. 3. SpyCatcher002–SpyTag002 conjugation positive control.** SDS-PAGE analysis of the reaction between free SpyCatcher002-CLP and free SpyTag002-MBP-CLP at defined time points (1, 5, 15, 30, and 60 min) at room temperature. The upper, middle, and lower bands correspond to the conjugation product, unreacted SpyTag002-MBP-CLP, and unreacted SpyCatcher002-CLP, respectively. The conjugation product band at 60 min was used as a reference standard representing 100% conjugation for densitometric quantification. M, molecular weight marker.

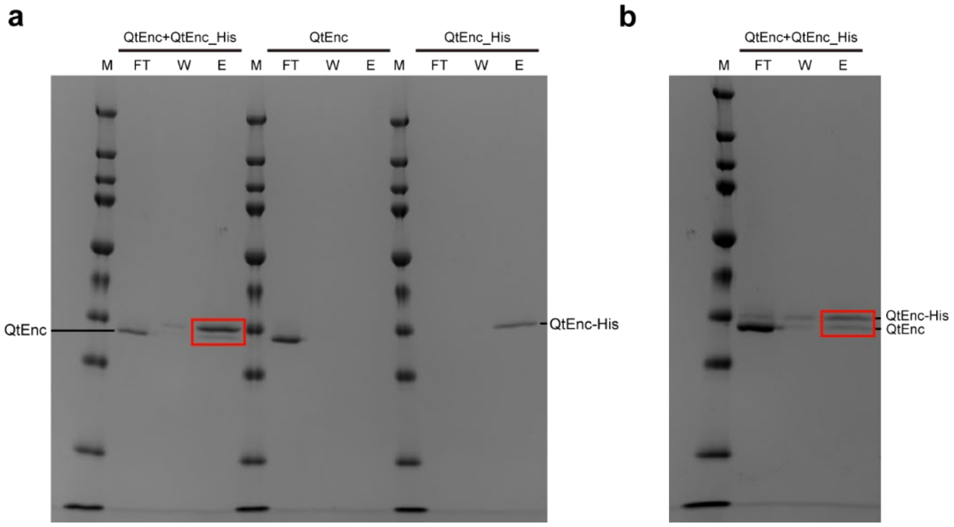


**Supplementary Fig. 4. Ni-NTA-based dynamic subunit-exchange assay of QtEnc and QtEnc-His.** A C-terminally His-tagged QtEnc variant (QtEnc-His) was mixed with untagged QtEnc for 30 min at 4°C, and the mixture was applied to a Ni-NTA spin column; QtEnc alone and QtEnc-His alone were analyzed in parallel as controls. **a**, For each sample (QtEnc + QtEnc-His, QtEnc alone, and QtEnc-His alone), the flow-through (FT), wash (W), and elution (E) fractions were analyzed by SDS-PAGE. Untagged QtEnc alone was recovered entirely in the flow-through with no signal in the elution, confirming that it has no intrinsic affinity for the column, whereas QtEnc-His alone was fully captured (no signal in the flow-through) and recovered in the elution. In the QtEnc + QtEnc-His mixture, the large majority of untagged QtEnc appeared in the flow-through, with only a small fraction co-eluting with QtEnc-His in the elution (red box), indicating limited subunit exchange between shells. **b**, The same experiment performed at higher shell concentration to increase the frequency of shell–shell contact; the co-eluting untagged QtEnc fraction (red box) increased relative to **a**, while the majority of untagged QtEnc still appeared in the flow-through. Together, these results indicate that subunit exchange between QtEnc shells is a rare and possibly contact-dependent event rather than a consequence of free, steady-state subunit dissociation. M, molecular weight marker.

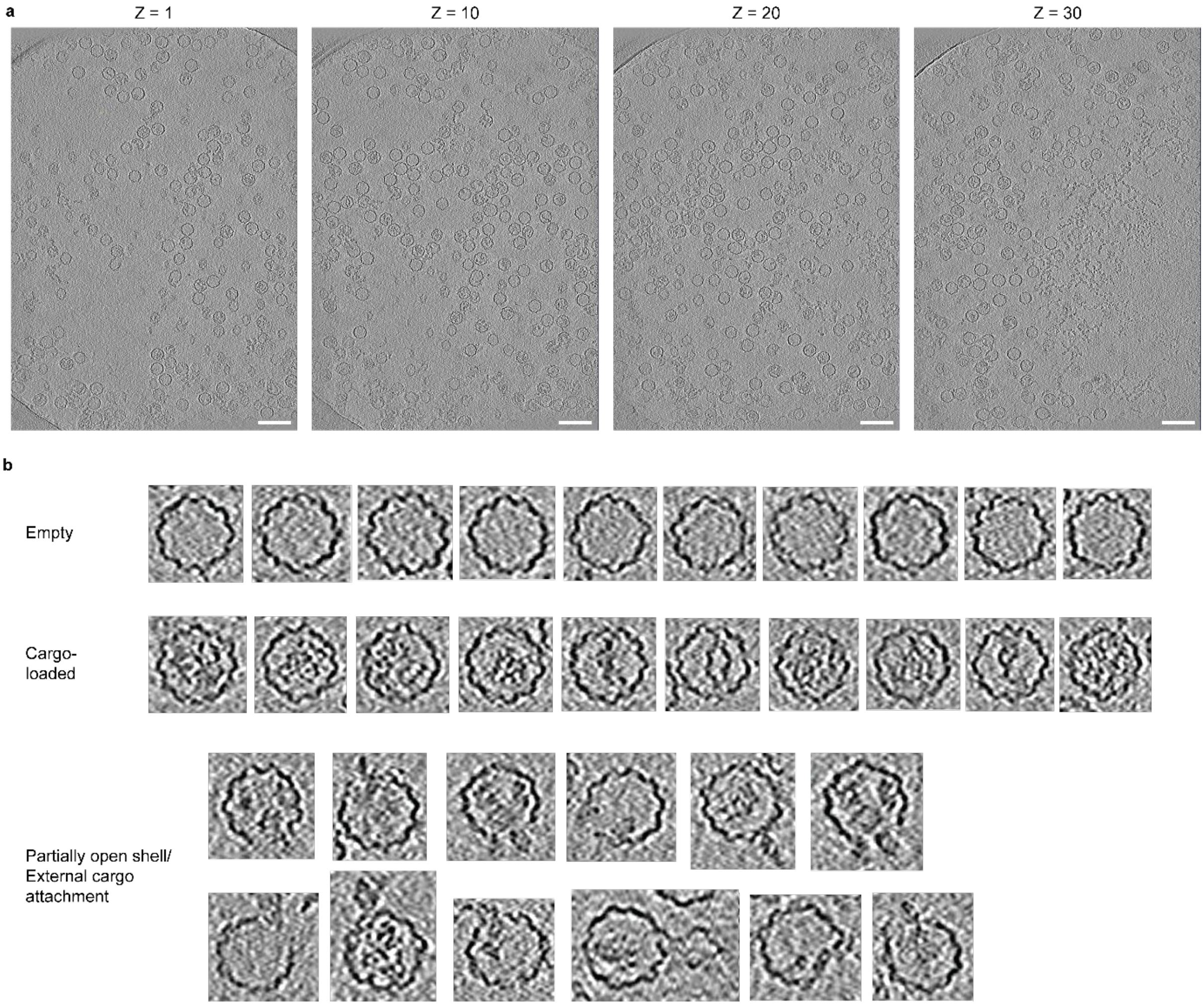


**Supplementary Fig. 5. Cryo-electron tomography of QtEnc mixed with β-galactosidase-CLP. a**, Representative z-slices (Z = 1, 10, 20, 30) through a reconstructed tomogram of QtEnc shortly after the addition of β-galactosidase-CLP, showing a field of intact, well-dispersed QtEnc shells with no indication of large-scale shell fragmentation or disassembly. **b**, Representative individual QtEnc shells extracted from the tomograms, grouped into three categories: empty shells (top), cargo-loaded shells containing internal β-galactosidase density (middle), and shells in a partially open state with some engaging cargo-like densities at the site of opening (bottom). The latter class is consistent with a transient loading intermediate in which cargo engages a locally opened shell element prior to internalization. Shells remained intact across all classes, with no evidence of global disassembly. Scale bars, 100 nm.

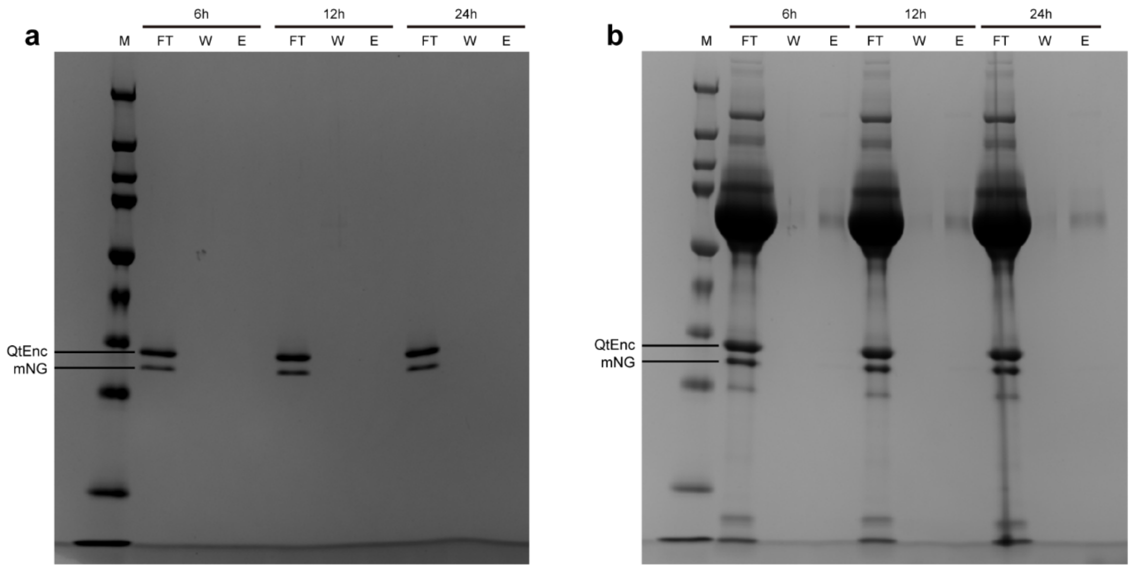


**Supplementary Fig. 6. Ni-NTA-based cargo retention assay of mNeonGreen-loaded QtEnc over time.** QtEnc loaded with His-tagged mNeonGreen was incubated in **a**, buffer (TBS) or **b**, supplemented cell culture medium (DMEM + 10% FBS + 1% PS) at 4°C for 6, 12, and 24 h, then applied to a Ni-NTA spin column. Released cargo, which exposes its His-tag, is captured on the column and recovered in the elution fraction, whereas encapsulated cargo, with its His-tag occluded by the shell, appears in the flow-through. For each time point, the flow-through (FT), wash (W), and elution (E) fractions were analyzed by SDS-PAGE. mNeonGreen and QtEnc was detected predominantly in the flow-through fractions, with no detectable mNeonGreen in the elution fractions at any time point, indicating that internalized cargo remains stably encapsulated for at least 24 h under both conditions. The strong high-molecular-weight bands in **b** correspond to serum proteins from the FBS-supplemented medium. M, molecular weight marker.

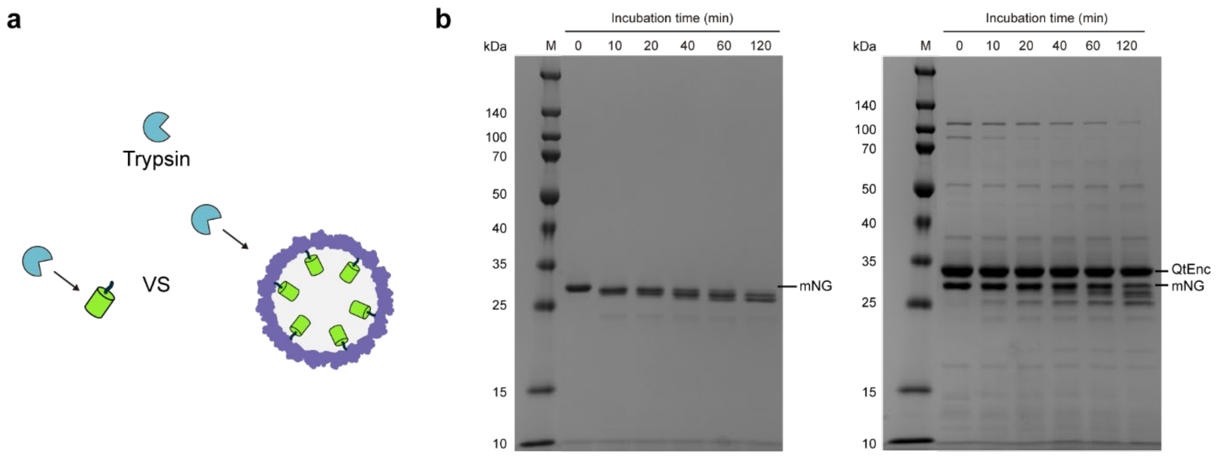


**Supplementary Fig. 7. Protease protection of encapsulated mNeonGreen by the QtEnc shell. a**, Schematic illustrating the trypsin proteolysis assay comparing free mNeonGreen (mNG) and QtEnc-loaded mNG. **b**, SDS-PAGE analysis of free mNG (left) and QtEnc-loaded mNG (right) incubated with trypsin at 37°C. Aliquots were taken at 0, 10, 20, 40, 60, and 120 min and subjected to SDS-PAGE analysis. Free mNG is completely cleaved within 10 min, with no intact full-length protein detectable. In contrast, full-length mNG remains detectable even after 120 min of incubation when encapsulated within QtEnc. M, molecular weight marker.

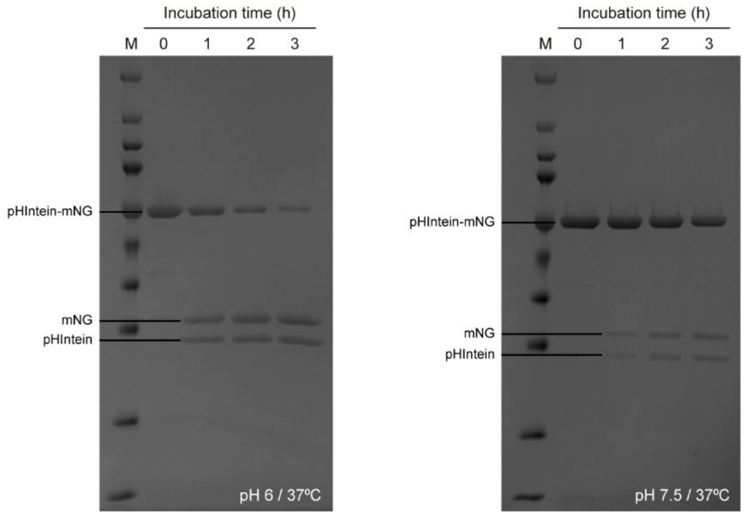


**Supplementary Fig. 8. pH-dependent cleavage activity of pHIntein.** SDS-PAGE analysis of CLP-pHIntein-mNeonGreen incubated at **a**, pH 6.0 and **b**, pH 7.5, both at 37°C. Aliquots were withdrawn at 0, 1, 2, and 3 h and subjected to SDS-PAGE analysis. The upper, middle, and lower bands correspond to the uncleaved CLP-pHIntein-mNeonGreen, cleaved mNeonGreen, and cleaved CLP-pHIntein fragments, respectively. At pH 6.0, progressive cleavage is observed over time, reaching approximately 85% cleavage efficiency within 3 h. At pH 7.5, cleavage is significantly attenuated, with only approximately 28% cleavage observed after 3 h, consistent with the reported pH-sensitive cleavage activity of pHIntein. M, molecular weight marker.

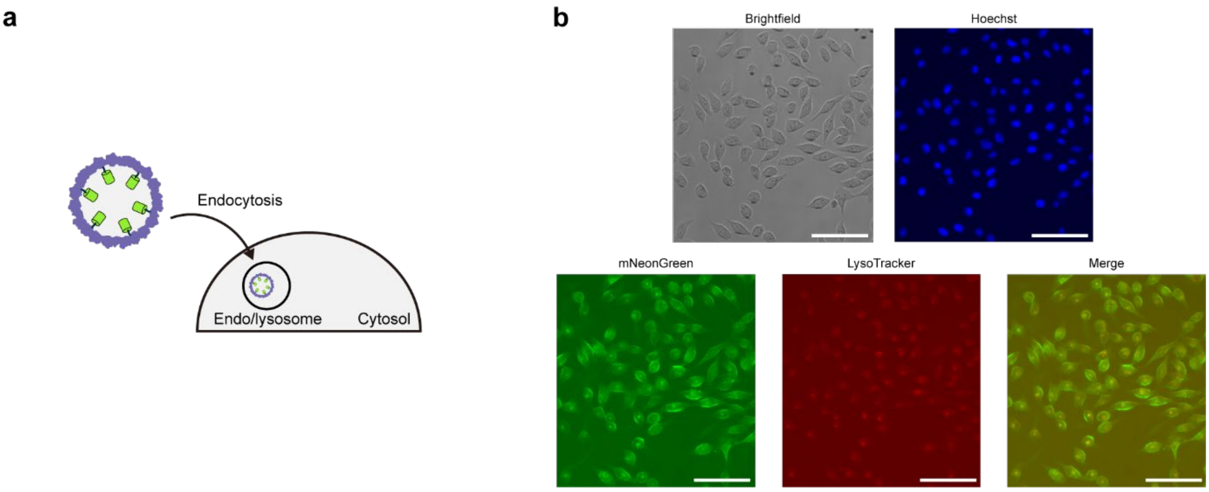


**Supplementary Fig. 9. Intracellular trafficking of mNeonGreen-loaded QtEnc in HeLa cells. a**, Schematic illustrating endocytic uptake of mNeonGreen-loaded QtEnc and its trafficking to acidic endo/lysosomal compartments. **b**, Representative fluorescence images of HeLa cells incubated with mNeonGreen-loaded QtEnc for 6 h, then washed to remove non-internalized material and stained with Hoechst (nuclei) and LysoTracker (acidic endo/lysosomal compartments). Panels show brightfield, Hoechst, mNeonGreen (QtEnc cargo), LysoTracker, and merged mNeonGreen/LysoTracker channels. Colocalization of internalized mNeonGreen with LysoTracker (merge) indicates that QtEnc is endocytosed and trafficked to acidic intracellular compartments. Images were acquired on an EVOS M5000 imaging system. Scale bars, 100 μm.

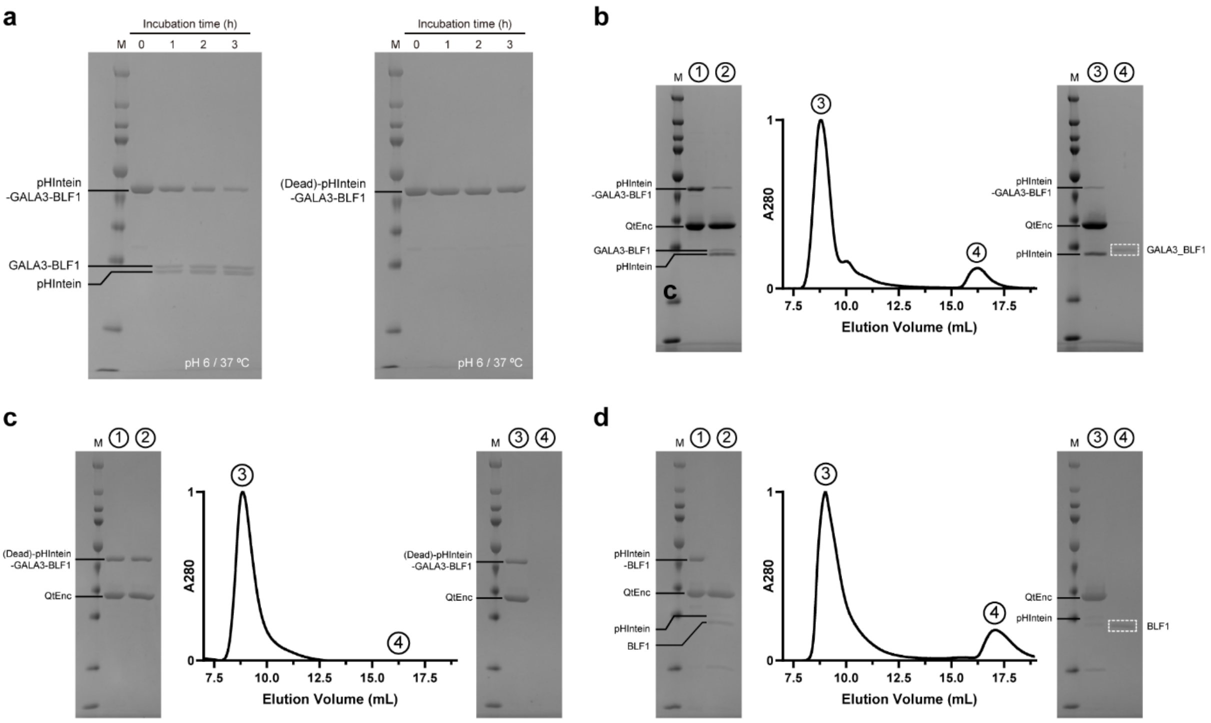


**Supplementary Fig. 10. Characterization of pHIntein-mediated cargo detachment and shell release for QtEncNC cargo constructs. a**, SDS-PAGE analysis of CLP-pHIntein-GALA3-BLF1 (left) and CLP-(Dead)-pHIntein-GALA3-BLF1 (right) incubated at pH 6.0 and 37°C. Aliquots were withdrawn at 0, 1, 2, and 3 h and subjected to SDS-PAGE analysis. Progressive cleavage is observed for CLP-pHIntein-GALA3-BLF1, whereas CLP-(Dead)-pHIntein-GALA3-BLF1 remains uncleaved, confirming the catalytic inactivity of (Dead)-pHIntein. **b-d**, SEC profiles and corresponding SDS-PAGE analyses of the pH-triggered POI release assay performed with QtEnc loaded with b, CLP-pHIntein-GALA3-BLF1, c, CLP-(Dead)-pHIntein-GALA3-BLF1, and d, CLP-pHIntein-BLF1. Numbered circles in the SEC profiles correspond to the annotated lanes in the SDS-PAGE gels: ① pre-acidification input; ② post-acidification pre-SEC sample; ③ (QtEnc-containing) void volume fraction; ④ released POI fraction. Quantitative release of b, GALA3-BLF1 and d, BLF1 from the QtEnc shell is observed upon pHIntein cleavage, whereas no POI release is detected for the c, (Dead)-pHIntein control. M, molecular weight marker.

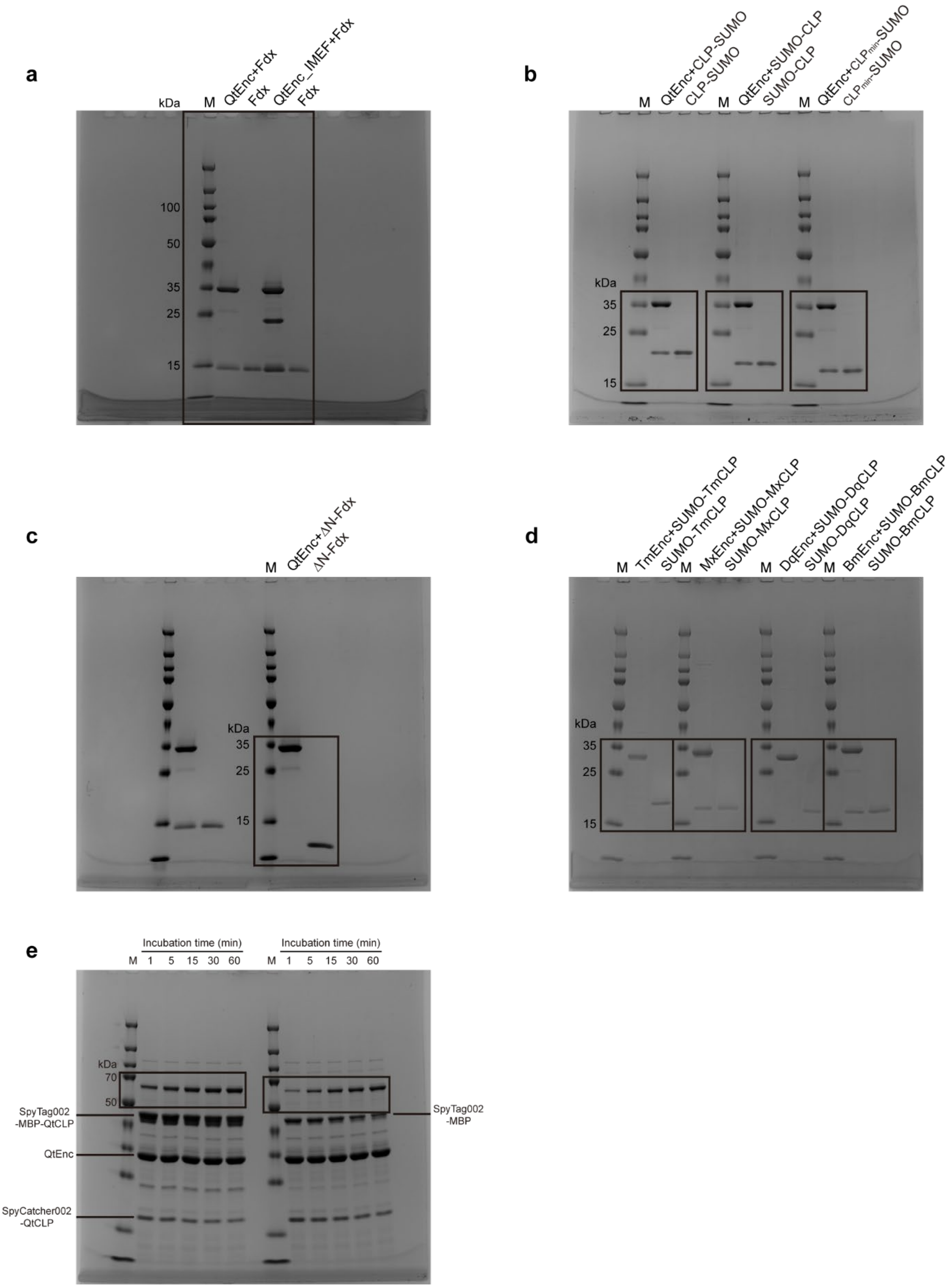
a
kDa
M
QtEnc+Fdx
Fdx
QtEnc_IMEF+Fdx
Fdx
100
50
35
25
15
b
M
QtEnc+CLP-SUMO
CLP-SUMO
M
QtEnc+SUMO-CLP
SUMO-CLP
M
QtEnc+CLPmin-SUMO
CLPmin-SUMO
kDa
35
25
15
c
M
QtEnc+ΔN-Fdx
ΔN-Fdx
kDa
35
25
15
d
M
TmEnc+SUMO-TmCLP
SUMO-TmCLP
M
MxEnc+SUMO-MxCLP
SUMO-MxCLP
M
DqEnc+SUMO-DqCLP
SUMO-DqCLP
M
BmEnc+SUMO-BmCLP
SUMO-BmCLP
kDa
35
25
15
e
Incubation time (min)
M 1 5 15 30 60
Incubation time (min)
M 1 5 15 30 60
kDa
70
50
SpyTag002
-MBP-QtCLP
QtEnc
SpyCatcher002
-QtCLP
SpyTag002
-MBP

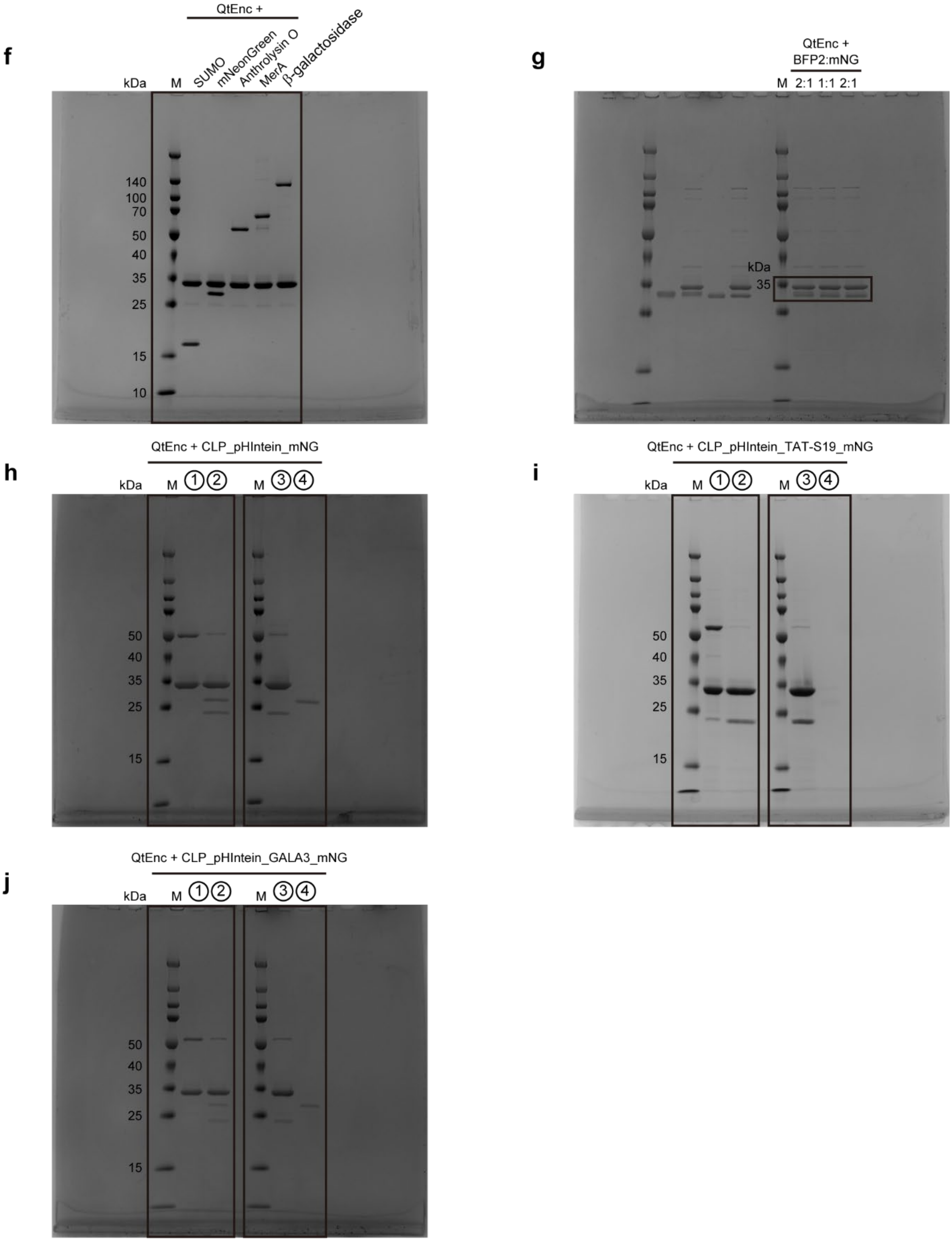
f
QtEnc +
kDa
M
SUMO
mNeonGreen
Anthrolysin O
MerA
β-galactosidase
140
100
70
50
40
35
25
15
10
g
QtEnc +
BFP2:mNG
M
2:1 1:1 2:1
kDa
35
h
QtEnc + CLP_pHIntein_mNG
kDa
M ①②
M ③④
50
40
35
25
15
i
QtEnc + CLP_pHIntein_TAT-S19_mNG
kDa
M ①②
M ③④
50
40
35
25
15
j
QtEnc + CLP_pHIntein_GALA3_mNG
kDa
M ①②
M ③④
50
40
35
25
15

**k**

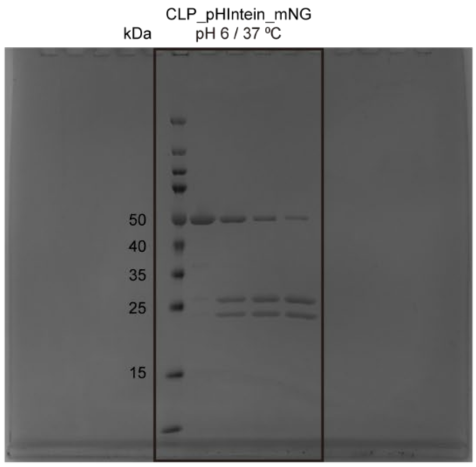
CLP_pHIntein_mNG
kDa
pH 6 / 37 ºC
50
40
35
25
15

**l**

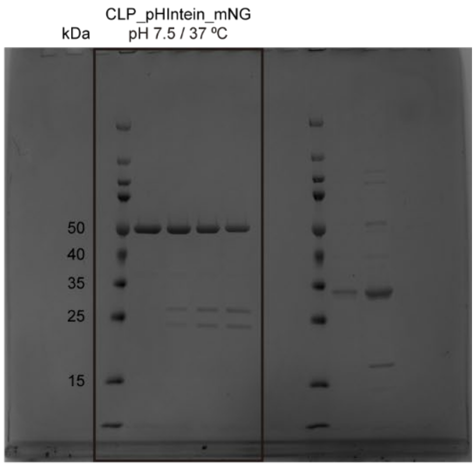
CLP_pHIntein_mNG
kDa
pH 7.5 / 37 ºC
50
40
35
25
15

**m**

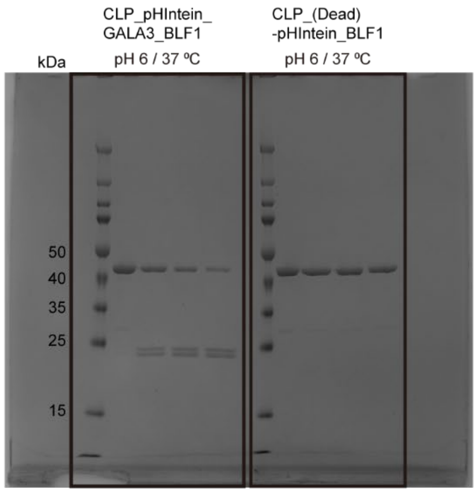
CLP_pHIntein_
GALA3_BLF1
CLP_(Dead)
-pHIntein_BLF1
kDa
pH 6 / 37 ºC
pH 6 / 37 ºC
50
40
35
25
15

**n**

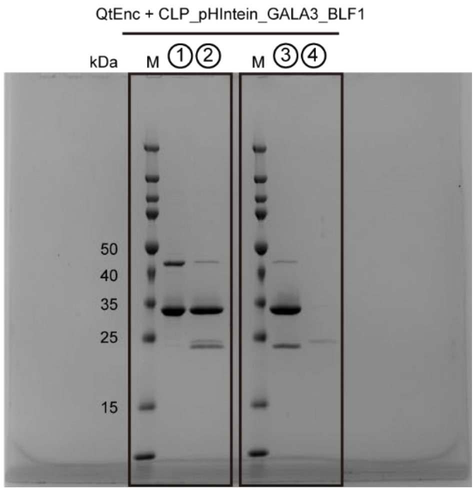
QtEnc + CLP_pHIntein_GALA3_BLF1
kDa
M ① ②
M ③ ④
50
40
35
25
15

**o**

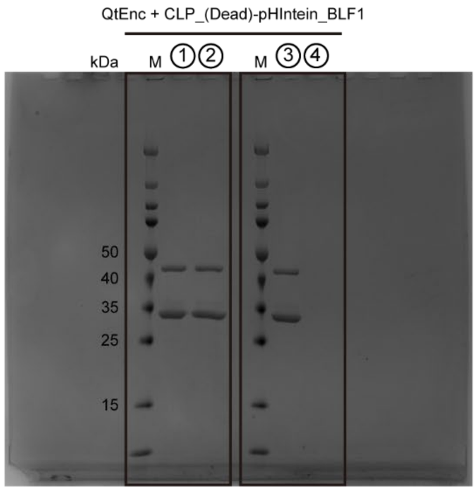
QtEnc + CLP_(Dead)-pHIntein_BLF1
kDa
M ① ②
M ③ ④
50
40
35
25
15

**p**

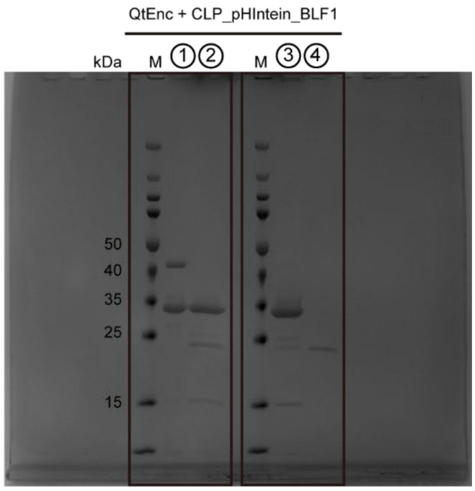
QtEnc + CLP_pHIntein_BLF1
kDa
M ① ②
M ③ ④
50
40
35
25
15

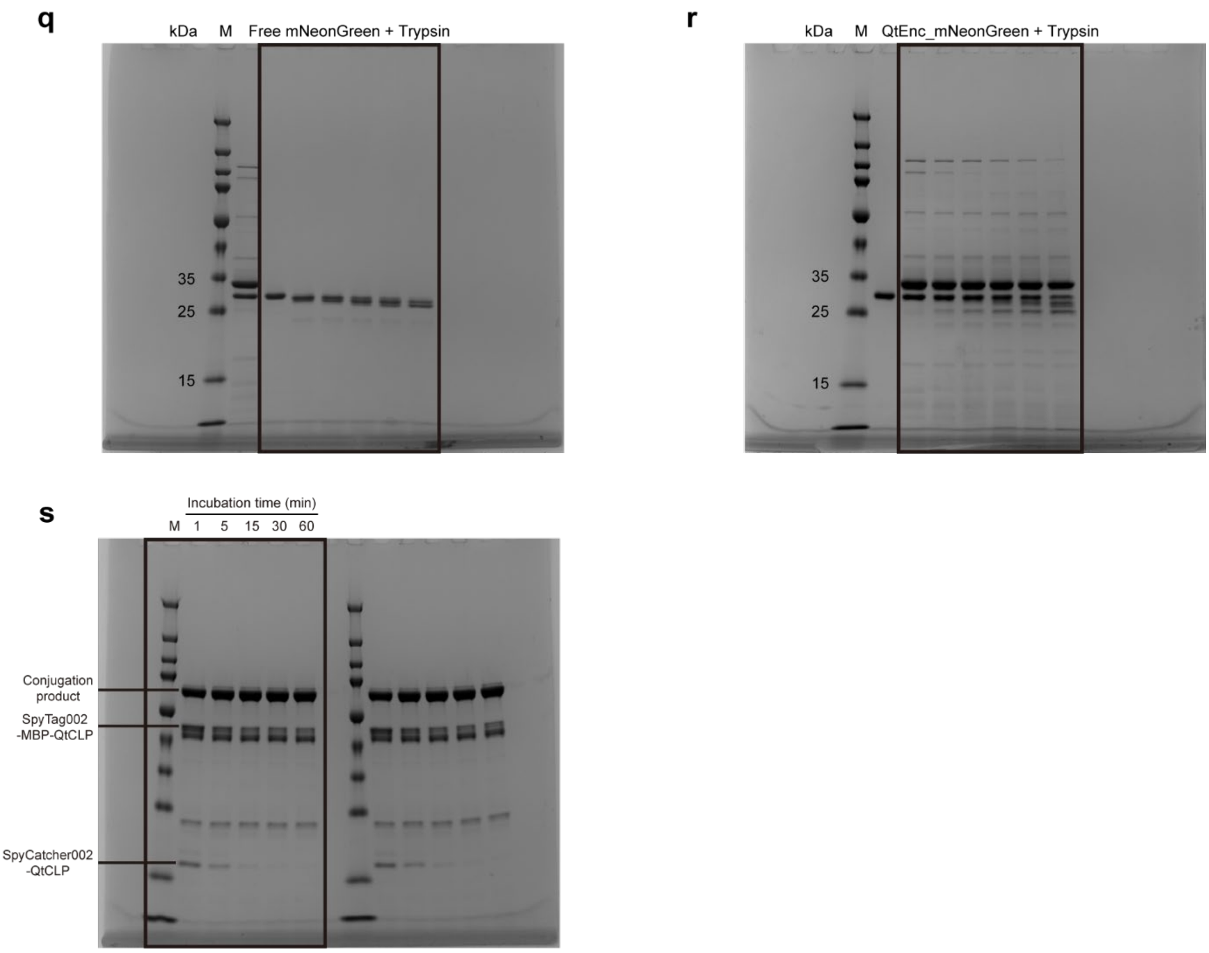


**Supplementary Fig. 11. Uncropped and annotated SDS-PAGE gels.** Uncropped versions of all SDS-PAGE gels presented in the main/SI figures. Boxes indicate the regions shown in the corresponding cropped figure panels. **a**, Fig. 1d. **b**, Fig. 2a. **c**, Fig. 2a. **d**, Fig. 2b. **e**, Fig. 2d. **f**, Fig. 3c. **g**, Fig. 4b. **h**, Fig. 5c. **i**, Fig. 6b. **j**, Fig. 6c. **k**, Supplementary Fig. 8. **l**, Supplementary Fig. 8. **m**, Supplementary Fig. 10a. **n**, Supplementary Fig. 10b. **o**, Supplementary Fig. 10c. **p**, Supplementary Fig. 10d. **q**, Supplementary Fig. 7b. **r**, Supplementary Fig. 7b. **s**, Supplementary Fig. 3.

**Supplementary Table 1. Amino acid sequences of proteins used in this study.**

| | |
|---|---|
| QtEnc | MNKSQLYPDSPLTDQDFNQLDQTVIEAARRQLVGRRFIELY GPLGRGMQSVFNDIFMESHEAKMDFQGSFDTEVESSRRVN YTIPMLYKDFVLYWRDLEQSKALDIPIDFSVAANAARDVAFLE DQMIFHGSKEFDIPGLMNVKGRLTHLIGNWYESGNAFQDIVE ARNKLLEMNHNGPYALVLSPELYSLLHRVHKDTNVLEIEHVR ELITAGVFQSPVLKGKSGVIVNTGRNNLDLAISEDFETAYLGE EGMNHPFRVYETVVLRIKRPAAICTLIDPEE |
| BmEnc | MGKTQLFPDSPLTDQDFSQLDQTVIDTARRQLIGRRFIELYG PLGRGIQSIFNDVFIENYEAKMDFQGSFDTDIETSKRVNYTIP LLYKDFVLYWRDLEQAKVLDIPIDFSAAANAARDVAILEDQMI FYGSKEFDIPGLMNVKGRSTHLIGNWYESGNAFQDVVEARN KLLEMKHNGPFALVLSPELYSLLHRVHKDTNVLEIEHVRELV TDGVFQTPVLKGKTGVLVNTGRNNLDLAVSEDFDTAYLGEE GMNHPFRVYETVVLRIKRPSAICTLEDGGE |
| DqEnc | MDFLDRNAAPLNEGEWQRIDEAVVSTARRTLVARRIIDVLGP LGSGVYSIPYSVFSGKSPTGIDMVGENEEFVVEASRRATINL PILYKDFKIMWRDVEADRHLGLPIDVSTAAVAGNFVAVQEDH LIFNGNVELGHDGLFTVKDRQTVAISDWNETGAALADVVKA VGALSQAGHYGPYAMVVSPVLFGRMIRVFGNTGMLELDQV KALITGGVHYSNVIGGSKAVVVATGSQNLNLALGQDMVTAY MGPTSMNHVFRVLETAALLVRRPDAICTIE |
| MxEnc | MPDFLGHAENPLREEEWARLNETVIQVARRSLVGRRILDIYG PLGAGVQTVPYDEFQGVSPGAVDIVGEQETAMVFTDARKFK TIPIIYKDFLLHWRDIEAARTHNMPLDVSAAAGAAALCAQQE DELIFYGDARLGYEGLMTANGRLTVPLGDWTSPGGGFQAIV EATRKLNEQGHFGPYAVVLSPRLYSQLHRIYEKTGVLEIETIR QLASDGVYQSNRLRGESGVVVSTGRENMDLAVSMDMVAA YLGASRMNHPFRVLEALLLRIKHPDAICTLEGAGATERR |
| TmEnc | MEFLKRSFAPLTEKQWQEIDNRAREIFKTQLYGRKFVDVEG PYGWEYAAHPLGEVEVLSDENEVVKWGLRKSLPLIELRATF TLDLWELDNLERGKPNVDLSSLEETVRKVAEFEDEVIFRGCE KSGVKGLLSFEERKIECGSTPKDLLEAIVRALSIFSKDGIEGP YTLVINTDRWINFLKEEAGHYPLEKRVEECLRGGKIITTPRIE DALVVSERGGDFKLILGQDLSIGYEDREKDAVRLFITETFTFQ VVNPEALILLKF |
| Qt-Fdx-His | MSHDRRLTVGSLLPNQPRPVAVPKAPSVVQPSKQPLPEKAII RLEQNGRSFSVRHVKGNLLTEGLSQGLPLQYKCRKGTCGV CTVKVTDGASRLSLPNQQEHKKLQGNIKSGFRLACQANME HHHHHH |
| Qt-ΔN-Fdx-His | MAIIRLEQNGRSFSVRHVKGNLLTEGLSQGLPLQYKCRKGT CGVCTVKVTDGASRLSLPNQQEHKKLQGNIKSGFRLACQAN MEHHHHHH |
| Qt-IMEF | MKEELDAFHQIFTTTKEAIERFMAMLTPVIENAEDDHERLYY HHIYEEEEQRLSRLDVLIPLIEKFQDETDEGLFSPSNNAFNRL LQELNLEKFGLHNFIEHVDLALFSFTDEERQTLLKELRKDAYE GYQYVKEKLAEINARFDHDYADPHAHHDEHRDHLADMPSA GSSHEEVQPVAHKKKGFTVGSLIQ |
| QtTP(N)-SUMO_His | MSHDRRLTVGSLLPNQPRPVAVPKAPSVVQPSKQPLPEKS DQEAKPSTEDLGDKKEGEYIKLKVIGQDSSEIHFKVKMTTHL KKLKESYCQRQGVPMNSLRFLFEGQRIADNHTPKELGMEEE DVIEVYQEQTGGSHHHHHH |
| Truncated-QtTP(N)-SUMO-His | MSHDRRLTVGSLLPGGSGGSSDQEAKPSTEDLGDKKEGEY IKLKVIGQDSSEIHFKVKMTTHLKKLKESYCQRQGVPMNSLR FLFEGQRIADNHTPKELGMEEEDVIEVYQEQTGGSHHHHHH |
| His-SUMO-QtTP(C) | MHHHHHHGGSSDQEAKPSTEDLGDKKEGEYIKLKVIGQDSS EIHFKVKMTTHLKKLKESYCQRQGVPMNSLRFLFEGQRIAD |

| | |
|---|---|
| | NHTPKELGMEEEDVIEVYQEQTGGSGGSHKKKGFTVGSLIQ |
| His-SUMO-DqTP(C) | MHHHHHHGGSSDQEAKPSTEDLGDKKEGEYIKLKVIGQDSS<br>EIHFKVKMTTHLKKLKESYCQRQGVPMNSLRFLFEGQRIAD<br>NHTPKELGMEEEDVIEVYQEQTGGSGGSDRIPTVGSMFGQ<br>K |
| His-SUMO-BmTP(C) | MHHHHHHGGSSDQEAKPSTEDLGDKKEGEYIKLKVIGQDSS<br>EIHFKVKMTTHLKKLKESYCQRQGVPMNSLRFLFEGQRIAD<br>NHTPKELGMEEEDVIEVYQEQTGGSGGSNKKKGFTVGSLI |
| His-SUMO-MxTP(C) | MHHHHHHGGSSDQEAKPSTEDLGDKKEGEYIKLKVIGQDSS<br>EIHFKVKMTTHLKKLKESYCQRQGVPMNSLRFLFEGQRIAD<br>NHTPKELGMEEEDVIEVYQEQTGGSGGSPEKRLTVGSLRR |
| His-SUMO-TmTP(C) | MHHHHHHGGSSDQEAKPSTEDLGDKKEGEYIKLKVIGQDSS<br>EIHFKVKMTTHLKKLKESYCQRQGVPMNSLRFLFEGQRIAD<br>NHTPKELGMEEEDVIEVYQEQTGGSGGSENTGGDLGIRKL |
| His-mNEON-QtTP(C) | MHHHHHHGGSVSKGEEDNMASLPATHELHIFGSINGVDFD<br>MVGQGTGNPNDGYEELNLKSTKGDLQFSPWILVPHIGYGFH<br>QYLPYPDGMSPFQAAMVDGSGYQVHRTMQFEDGASLTVN<br>YRYTYEGSHIKGEAQVKGTGFPADGPVMTNSLTAADWCRS<br>KKTYPNDKTIISTFKWSYTTGNGKRYRSTARTTYTFAKPMAA<br>NYLKNQPMYVFRKTELKHSKTELNFKEWQKAFTDVMGMDE<br>LYKGGSGGSHKKKGFTVGSLIQ |
| His-mTagBFP2-QtTP(C) | MHHHHHHGGSVSKGEELIKENMHMKLYMEGTVDNHHFKCT<br>SEGEGKPYEGTQTMRIKVVEGGPLPFAFDILATSFLYGSKTFI<br>NHTQGIPDFFKQSFPEGFTWERVTTYEDGGVLTATQDTSLQ<br>DGCLIYNVKIRGVNFTSNGPVMQKKTLGWEAFTETLYPADG<br>GLEGRNDMALKLVGGSHLIANAKTTYRSKKPAKNLKMPGVY<br>YVDYRLERIKEANNETYVEQHEVAVARYCDLPSKLGHKLNG<br>GSGGSHKKKGFTVGSLIQ |
| His-mCherry-QtTP(C) | MHHHHHHGGSVSKGEEDNMAIIKEFMRFKVHMEGSVNGHE<br>FEIEGEGEGRPYEGTQTAKLKVTKGGPLPFAWDILSPQFMY<br>GSKAYVKHPADIPDYLKLSFPEGFKWERVMNFEDGGVVTVT<br>QDSSLQDGEFIYKVKLRGTNFPSDGPVMQKKTMGWEASSE<br>RMYPEDGALKGEIKQRLKLKDGGHYDAEVKTTYKAKKPVQL<br>PGAYNVNIKLDITSHNEDYTIVEQYERAEGRHSTGGMDELYK<br>GGSGGSHKKKGFTVGSLIQ |
| His-spycatcher002-QtTP(C) | MSYYHHHHHHDYDIPTTENLYFQGAMVTTLSGLSGEQGPS<br>GDMTTEEDSATHIKFSKRDEDGRELAGATMELRDSSGKTIS<br>TWISDGHVKDFYLYPGKYTFVETAAPDGYEVATAITFTVNEQ<br>GQVTVNGEATKGDAHTGSSGSGGSGGSHKKKGFTVGSLIQ |
| His-SpyTag002-MBP-QtTP(C) | MGSSHHHHHHSSGLVPRGSVPTIVMVDAYKRYKGSGESGK<br>IEEGKLVIWINGDKGYNGLAEVGKKFEKDTGIKVTVEHPDKL<br>EEKFPQVAATGDGPDIIFWAHDRFGGYAQSGLLAEITPDKAF<br>QDKLYPFTWDAVRYNGKLIAYPIAVEALSLIYNKDLLPNPPKT<br>WEEIPALDKELKAKGKSALMFNLQEPYFTWPLIAADGGYAFK<br>YENGKYDIKDVGVDNAGAKAGLTFLVDLIKNKHMNADTDYSI<br>AEAAFNKGETAMTINGPWAWSNIDTSKVNYGVTVLPTFKGQ<br>PSKPFVGVLSAGINAASPNKELAKEFLENYLLTDEGLEAVNK<br>DKPLGAVALKSYEEELAKDPRIAATMENAQKGEIMPNIPQMS<br>AFWYAVRTAVINAASGRQTVDEALKDAQTNSSSGGSGGSH<br>KKKGFTVGSLIQ |
| His-AnthrolysinO-QtTP(C) | MHHHHHHAAAMETQAGNATGAIKNASDINTGIANLKYDSRDI<br>LAVNGDKVESFIPKESINSNGKFVVVEREKKSLTTSPVDILIID<br>SVVNRTYPGAVQLANKAFADNQPSLLVAKRKPLNISIDLPGM<br>RKENTITVQNPTYGNVAGAVDDLVSTWNEKYSTTHTLPARM<br>QYTESMVYSKSQIASALNVNAKYLDNSLNIDFNAVANGEKKV<br>MVAAYKQIFYTVSAELPNNPSDLFDNSVTFDELTRKGVSNSA<br>PPVMSVSNVAYGRTVYVKLETTSKSKDVQAAFKALLKNNSVE |

| | |
|---|---|
| | TSGQYKDIFEESTFTAVVLGGDAKEHNKVVTKDFNEIRNIIKD<br>NAELSFKNPAYPISYTSTFLKDNATAAVHNNTDYIETTTTEYS<br>SAKMTLDHYGAYVAQFDVSWDEFTFDQNGKEVLTHKTWEG<br>SGKDKTAHYSTVIPLPPNSKNIKIVARECTGLAWEWWRTIINE<br>QNVPLTNEIKVSIGGTTLYPTATISHGGGSGGGSGGGSHKK<br>KGFTVGSLIQ |
| His-TEV-Truncated_QtTP(N)-MerA | MHHHHHHHHGGGSGGGSENLYFQGMSHDRRLTVGSLLPG<br>GSGGSTHLKITGMTCDSCAAHVKEALEKVPGVQSALVSYPK<br>GTAQLAIVPGTSPDALTAAVAGLGYKATLADAPLADNRVGLL<br>DKVRGWMAAAEKHSGNEPPVQVAVIGSGGAAMAAALKAVE<br>QGAQVTLIERGTIGGTCVNVGCVPSKIMIRAAHIAHLRRESPF<br>DGGIAATVPTIDRSKLLAQQQARVDELRHAKYEGILGGNPAI<br>TVVHGEARFKDDQSLTVRLNEGGERVVMFDRCLVATGASP<br>AVPPIPGLKESPYWTSTEALASDTIPERLAVIGSSVVALELAQ<br>AFARLGSKVTVLARNTLFFREDPAIGEAVTAAFRAEGIEVLEH<br>TQASQVAHMDGEFVLTTTHGELRADKLLVATGRTPNTRSLA<br>LDAAGVTVNAQGAIVIDQGMRTSNPNIYAAGDCTDQPQFVY<br>VAAAAGTRAAINMTGGDAALDLTAMPAVVFTDPQVATVGYS<br>EAEAHHDGIETDSRTLTLDNVPRALANFDTRGFIKLVIEEGSH<br>RLIGVQAVAPEAGELIQTAALAIRNRMTVQELADQLFPYLTM<br>VEGLKLAAQTFNKDVKQLSCCA |
| His-beta-galactosidase-QtTP(C) | MHHHHHHGGSGGSGGSMTMITDSLAVVLQRRDWENPGVT<br>QLNRLAAHPPFASWRNSEEARTDRPSQQLRSLNGEWRFA<br>WFPAPEAVPESWLECDLPEADTVVVPSNWQMHGYDAPIYT<br>NVTYPITVNPPFVPTENPTGCYSLTFNVDESWLQEGQTRIIF<br>DGVNSAFHLWCNGRWVGYGQDSRLPSEFDLSAFLRAGEN<br>RLAVMVLRWSDGSYLEDQDMWRMSGIFRDVSLLHKPTTQI<br>SDFHVATRFNDDFSRAVLEAEVQMCGELRDYLRVTVSLWQ<br>GETQVASGTAPFGGEIIDERGGYADRVTLRLNVENPKLWSA<br>EIPNLYRAVVELHTADGTLIEAEACDVGFREVRIENGLLLLNG<br>KPLLIRGVNRHEHHPLHGQVMDEQTMVQDILLMKQNNFNAV<br>RCSHYPNHPLWYTLCDRYGLYVVDEANIETHGMVPMNRLT<br>DDPRWLPAMSERVTRMVQRDRNHPSVIIWSLGNESGHGAN<br>HDALYRWIKSVDPSRPVQYEGGGADTTATDIICPMYARVDE<br>DQPFPAVPKWSIKKWLSLPGETRPLILCEYAHAMGNSLGGF<br>AKYWQAFRQYPRLQGGFVWDWVDQSLIKYDENGNPWSAY<br>GGDFGDTPNDRQFCMNGLVFADRTPHPALTEAKHQQQFFQ<br>FRLSGQTIEVTSEYLFRHSDNELLHWMVALDGKPLASGEVP<br>LDVAPQGKQLIELPELPQPESAGQLWLTVRVVQPNATAWSE<br>AGHISAWQQWRLAENLSVTLPAASHAIPHLTTSEMDFCIELG<br>NKRWQFNRQSGFLSQMWIGDKKQLLTPLRDQFTRAPLDND<br>IGVSEATRIDPNAWVERWKAAGHYQAEAALLQCTADTLADA<br>VLITTAHAWQHQGKTLFISRKTYRIDGSGQMAITVDVEVASD<br>TPHPARIGLNCQLAQVAERVNWLGLGPQENYPDRLTAACFD<br>RWDLPLSDMYTPYVFPSENGLRCGTRELNYGPHQWRGDF<br>QFNISRYSQQQLMETSHRHLLHAEEGTWLNIDGFHMGIGGD<br>DSWSPSVSAEFQLSAGRYHYQLVWCQKGGGSGGGSGGG<br>SGGGSHKKKGFTVGSLIQ |
| His-Truncated-QtTP(N)-pHIntein-mNEON | MHHHHHHGGGGSMSHDRRLTVGSLLPGGGSGGGGSGGG<br>GSGGGGSGALAEGTRIFDPVTGTTHRIEDVVGGRKPIHVVA<br>AAKDGTLHARPVVSWFDQGTRDVIGLRIAGGAILWATPDHK<br>VLTEYGWRAAGELRKGDRVAQPRRFDGFGDSAPIPARVQA<br>LADALDDKFLHDMLAEELRYSVIREVLPTRRARTFNLEVEEL<br>HTLVAEGVVVHNSMVSKGEEDNMASLPATHELHIFGSINGV<br>DFDMVGQGTGNPNDGYEELNLKSTKGDLQFSPWILVPHIGY<br>GFHQYLPYPDGMSPFQAAMVDGSGYQVHRTMQFEDGASL<br>TVNYRYTYEGSHIKGEAQVKGTGFPADGPVMTNSLTAADW<br>CRSKKTYPNDKTIISTFKWSYTTGNGKRYRSTARTTYTFAKP |

| | |
|---|---|
| | MAANYLKNQPMYVFRKTELKHSKTELNFKEWQKAFTDVMG MDELYK |
| His-Truncated-QtTP(N)-pHIntein-TAT-S19_mNEON | MHHHHHHGGGGSMSHDRRLTVGSLLPGGGSGGGGSGGG GSGGGGSGALAEGTRIFDPVTGTTHRIEDVVGGRKPIHVVA AAKDGTLHARPVVSWFDQGTRDVIGLRIAGGAILWATPDHK VLTEYGWRAAGELRKGDRVAQPRRFDGFGDSAPIPARVQA LADALDDKFLHDMLAEELRYSVIREVLPTRRARTFNLEVEEL HTLVAEGVVVHNSYGRKKRRQRRRPFVIGAGVLGALGTGIG GIGGGGSMVSKGEEDNMASLPATHELHIFGSINGVDFDMVG QGTGNPNDGYEELNLKSTKGDLQFSPWILVPHIGYGFHQYL PYPDGMSPFQAAMVDGSGYQVHRTMQFEDGASLTVNYRY TYEGSHIKGEAQVKGTGFPADGPVMTNSLTAADWCRSKKT YPNDKTIISTFKWSYTTGNGKRYRSTARTTYTFAKPMAANYL KNQPMYVFRKTELKHSKTELNFKEWQKAFTDVMGMDELYK |
| His-Truncated-QtTP(N)-pHIntein-GALA3-mNEON | MHHHHHHGGGGSMSHDRRLTVGSLLPGGGSGGGGSGGG GSGGGGSGALAEGTRIFDPVTGTTHRIEDVVGGRKPIHVVA AAKDGTLHARPVVSWFDQGTRDVIGLRIAGGAILWATPDHK VLTEYGWRAAGELRKGDRVAQPRRFDGFGDSAPIPARVQA LADALDDKFLHDMLAEELRYSVIREVLPTRRARTFNLEVEEL HTLVAEGVVVHNSLAEALAEALEALAAGGGSGGGSMVSKG EEDNMASLPATHELHIFGSINGVDFDMVGQGTGNPNDGYEE LNLKSTKGDLQFSPWILVPHIGYGFHQYLPYPDGMSPFQAA MVDGSGYQVHRTMQFEDGASLTVNYRYTYEGSHIKGEAQV KGTGFPADGPVMTNSLTAADWCRSKKTYPNDKTIISTFKWS YTTGNGKRYRSTARTTYTFAKPMAANYLKNQPMYVFRKTEL KHSKTELNFKEWQKAFTDVMGMDELYK |
| His-Truncated-QtTP(N)-pHIntein-GALA3-BLF1 | MHHHHHHGGGGSMSHDRRLTVGSLLPGGGSGGGGSGGG GSGGGGSGALAEGTRIFDPVTGTTHRIEDVVGGRKPIHVVA AAKDGTLHARPVVSWFDQGTRDVIGLRIAGGAILWATPDHK VLTEYGWRAAGELRKGDRVAQPRRFDGFGDSAPIPARVQA LADALDDKFLHDMLAEELRYSVIREVLPTRRARTFNLEVEEL HTLVAEGVVVHNSLAEALAEALEALAAGGGSGGGSMPNSLE AQIRQAMKTGSTLTIEFDQALNQKSPGTLNVFLHPANGGVRI DLDSGNQGEPAKILWLPWKQGELQTLQPGSISTVDMLFFTY YLSGCKVFAGDGGPVWHIDAPVEANQFWRRMSSDEWMED WEVGTDRQVAYLHRAGQSDSLWNLSAYLEGAAPSTYGRD NLGQAVVGGIVTGRQQMSLYQYATTSSGSSAWSPLTYTLQ QRKQ |
| His-Truncated-QtTP(N)-(Dead)-pHIntein-GALA3-BLF1 | MHHHHHHGGGGSMSHDRRLTVGSLLPGGGSGGGGSGGG GSGGGGSGALAEGTRIFDPVTGTTHRIEDVVGGRKPIHVVA AAKDGTLHARPVVSWFDQGTRDVIGLRIAGGAILWATPDHK VLTEYGWRAAGELRKGDRVAQPRRFDGFGDSAPIPARVQA LADALDDKFLHDMLAEELRYSVIREVLPTRRARTFNLEVEEL HTLVAEGVVVHASLAEALAEALEALAAGGGSGGGSMPNSLE AQIRQAMKTGSTLTIEFDQALNQKSPGTLNVFLHPANGGVRI DLDSGNQGEPAKILWLPWKQGELQTLQPGSISTVDMLFFTY YLSGCKVFAGDGGPVWHIDAPVEANQFWRRMSSDEWMED WEVGTDRQVAYLHRAGQSDSLWNLSAYLEGAAPSTYGRD NLGQAVVGGIVTGRQQMSLYQYATTSSGSSAWSPLTYTLQ QRKQ |
| His-Truncated-QtTP(N)-pHIntein-BLF1 | MHHHHHHGGGGSMSHDRRLTVGSLLPGGGSGGGGSGGG GSGGGGSGALAEGTRIFDPVTGTTHRIEDVVGGRKPIHVVA AAKDGTLHARPVVSWFDQGTRDVIGLRIAGGAILWATPDHK VLTEYGWRAAGELRKGDRVAQPRRFDGFGDSAPIPARVQA LADALDDKFLHDMLAEELRYSVIREVLPTRRARTFNLEVEEL HTLVAEGVVVHNSMPNSLEAQIRQAMKTGSTLTIEFDQALN QKSPGTLNVFLHPANGGVRIDLDSGNQGEPAKILWLPWKQ GELQTLQPGSISTVDMLFFTYYLSGCKVFAGDGGPVWHIDA |

| | |
|---|---|
| | PVEANQFWRRMSSDEWMEDWEVGTDRQVAYLHRAGQSD SLWNLSAYLEGAAPSTYGRDNLGQAVVGGIVTGRQQMSLY QYATTSSGSSAWSPLTYTLQQRKQ |

**Supplementary Table 2. DNA primers used in this study.**

| Primer | Sequence (5'->3') |
|---|---|
| pETDuet_F | ATTAACCTAGGCTGCTGC |
| pETDuet_B | TGTATATCTCCTTCTTATACTTAACTAATATAC |
| ΔN-Fdx_F | AAGTATAAGAAGGAGATATACAATGGCCATTATTCGTCTGG |
| ΔN-Fdx_B | GCAGCAGCCTAGGTTAATTC |
| $CLP_{min}$-SUMO_F | GGTAGGTTCCCTTTTACCAGGCGGTAGTGGCGGTTCGTCTGACCAAGAGGCGAAACC |
| $CLP_{min}$-SUMO_B | TGGTAAAAGGGAACCTACCG |
| CLP-(Dead)-pHIntein -GALA3-BLF1_F | GGAGGGTGTAGTTGTGCACGCGTCGCTTGCAGAGGCGCTGG |
| CLP-(Dead)-pHIntein -GALA3-BLF1_B | GTGCACAACTACACCCTCC |

**Supplementary Table 3. Cryo-EM data collection and refinement statistics.**

| | QtEnc_FDX (EMDB-EMD: 76203) (PDB ID: 11YX) |
|---|---|
| **Data collection and processing** | |
| Magnification | 105,000 |
| Voltage (kV) | 300 |
| Electron exposure (e–/Å$^2$) | 57.2 |
| Defocus range (μm) | -1.0 to -1.8 |
| Pixel size (Å) | 0.832 (downsampled to 1.09 Å) |
| Symmetry imposed | I |
| Initial particle images (no.) | 78,442 |
| Final particle images (no.) | 70,540 |
| Map resolution (Å) | 2.47 |
| FSC threshold | 0.143 |
| **Refinement** | |
| Initial model used (PDB code) | |
| Model resolution (Å) | 2.8 |
| FSC threshold | 0.5 |
| Map sharpening *B* factor (Å$^2$) | -83.7 |
| Model composition | |
| Non-hydrogen atoms | 9,201 |
| Protein residues | 1,145 |
| Ligands | 0 |
| *B* factors (Å$^2$) | |
| Protein | 35.69 |
| R.M.S. deviations | |
| Bond lengths (Å) | 0.005 |
| Bond angles (°) | 1.014 |
| Validation | |
| MolProbity score | 1.30 |
| Clashscore | 5.16 |
| Poor rotamers (%) | 0.3 |
| Ramachandran plot | |
| Favored (%) | 97.88 |
| Allowed (%) | 2.12 |
| Disallowed (%) | 0 |
